# ROR1 plays a critical role in pancreatic tumor-initiating cells with a partial EMT signature

**DOI:** 10.1101/2022.07.13.499868

**Authors:** Masaya Yamazaki, Shinjiro Hino, Shingo Usuki, Yoshihiro Miyazaki, Tatsuya Oda, Mitsuyoshi Nakao, Takaaki Ito, Kazuya Yamagata

## Abstract

Tumor-initiating cells are the major drivers of chemoresistance and relapse, making them attractive targets for cancer therapy. However, the identity of tumor-initiating cells in human pancreatic ductal adenocarcinoma (PDAC) and the key molecules underlying their traits remain poorly understood. Here, we show that a partial epithelial-mesenchymal transition (EMT)-like subpopulation marked by high expression of receptor tyrosine kinase-like orphan receptor 1 (ROR1) is the origin of heterogeneous tumor cells in PDAC. We demonstrate that ROR1 depletion suppresses tumor growth, recurrence after chemotherapy, and metastasis. Mechanistically, ROR1 induces the expression of AURKB by activating E2F to enhance PDAC proliferation. Furthermore, epigenomic analyses reveal that *ROR1* is transcriptionally dependent on YAP/BRD4 binding at the enhancer region, and targeting this pathway reduces *ROR1* expression and prevents PDAC growth. Collectively, our findings reveal a critical role of ROR1^high^ cells as tumor-initiating cells and the functional importance of ROR1 in PDAC progression, thereby highlighting its therapeutic targetability.

**Graphical abstract:** 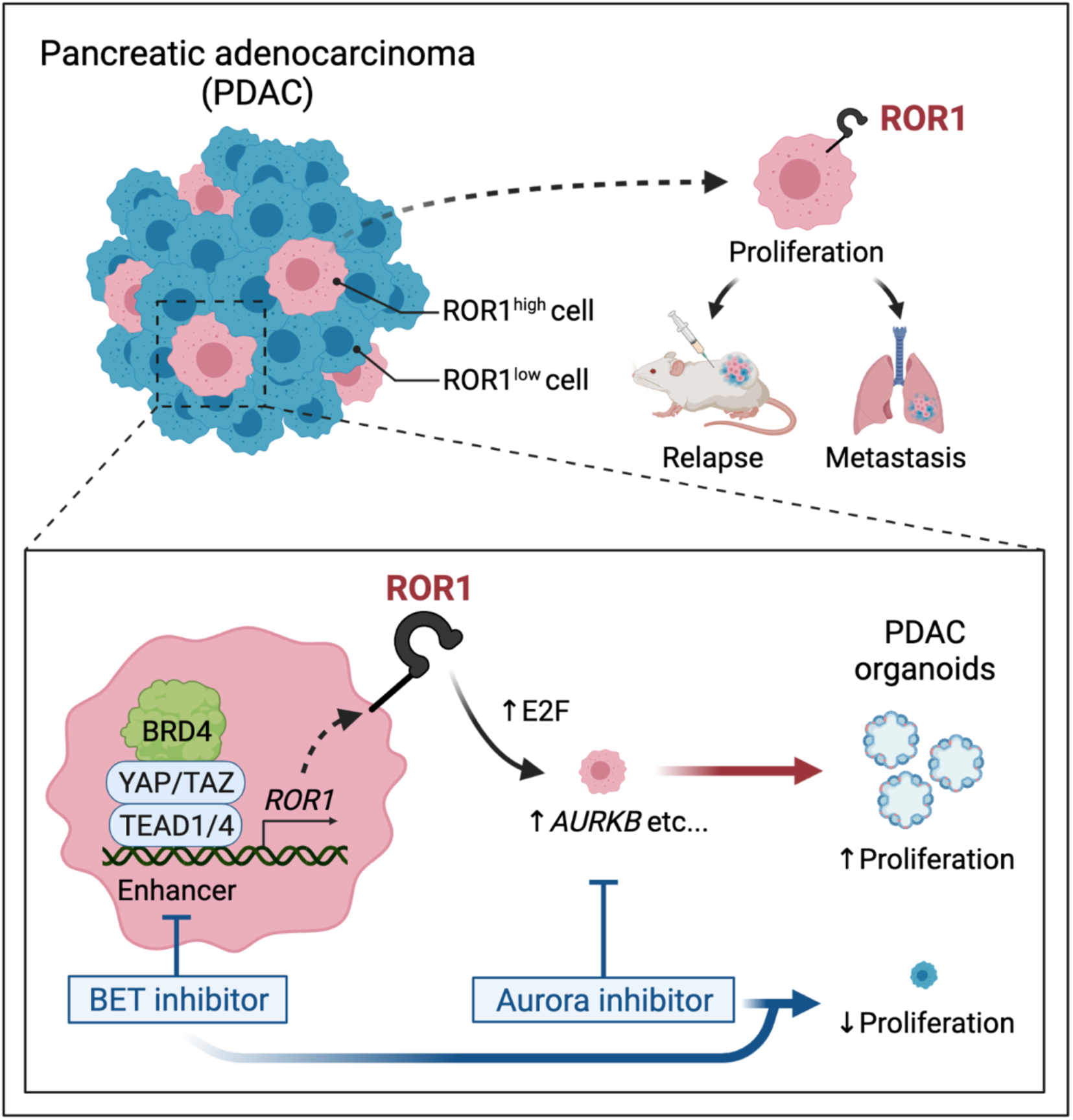

## Introduction

Pancreatic ductal adenocarcinoma (PDAC) is a malignant neoplasm with a poor prognosis. PDAC is highly aggressive, with a propensity for both local invasion and distant metastasis in the early stage, and is resistant to most treatments (Kleeff *et al*, 2016; Mizrahi *et al*, 2020), resulting in a 5-year overall survival rate of approximately 10% (National Cancer Institute. Cancer Stat Facts: Pancreatic Cancer. https://seer.cancer.gov/statfacts/html/pancreas.html). Deep whole-exome sequencing identified a high frequency of somatic DNA mutations in *KRAS* (>90%), *TP53* (>70%), *CDKN2A*, and *SMAD4* (both approximately 30%) (Raphael *et al*, 2017). Moreover, 60% of the tumors with wild-type *KRAS* harbored an alternative RAS-MAPK activating pathway, which highlights the importance of the RAS pathway in PDAC. Recently, inhibitors against mutant KRAS^G12D^ or KRAS^G12C^ have been developed and have given hope for breakthrough therapies in malignancies with *KRAS* mutations, including PDAC (Wang *et al*, 2022; Mao *et al*, 2022; Canon *et al*, 2019). However, resistance mechanisms have become uncovered, and thus, other therapeutic strategies have also been explored (Xue *et al*, 2020; Awad *et al*, 2021).

Intratumor heterogeneity contributes substantially to the characteristics of aggressive tumors, such as high frequencies of metastasis and resistance to treatments (Shibue & Weinberg, 2017). In particular, tumor-initiating cells (also referred to as cancer stem cells: CSCs) have been considered the source of cellular heterogeneity (Clarke *et al*, 2006; Shackleton *et al*, 2009; Magee *et al*, 2012). Their highly plastic nature allows tumor-initiating cells to generate a cellular hierarchy similar to that of normal tissue (Marusyk *et al*, 2020). The existence and importance of tumor-initiating cells are evidenced by their capability for tumorigenesis, metastasis, and recurrence in multiple malignancies (Batlle & Clevers, 2017; Bonnet & Dick, 1997; Al-Hajj *et al*, 2003; Singh *et al*, 2003; O’Brien *et al*, 2007; Ricci-Vitiani *et al*, 2007; Dalerba *et al*, 2007; Barker *et al*, 2009; De Sousa E Melo *et al*, 2017; Shimokawa *et al*, 2017). Thus, tumor-initiating cells play a central role in tumor progression. In PDAC, previous studies of intratumor heterogeneity have shown that subpopulations of cells marked by CD44^+^/CD24^+^/ESA^+^ (Li *et al*, 2007), CD133 (Hermann *et al*, 2007), DCLK1 (Bailey *et al*, 2014), and Musashi (Fox *et al*, 2016; Lytle *et al*, 2019) are functionally distinct tumor-initiating cells. However, at single-cell resolution, intratumor heterogeneity involved in disease progression is poorly understood. In addition, the therapeutic benefit of eradicating tumor-initiating cells in PDAC remains unclear. To identify targetable tumor-initiating cells, we here conducted single-cell RNA sequencing and functional approaches using human PDAC xenografts. We found that a partial epithelial-mesenchymal transition (EMT)-like subpopulation with high expression of receptor tyrosine kinase-like orphan receptor 1 (ROR1) serves as a source of intratumor heterogeneity. We demonstrated that ROR1^high^ cells have high tumorigenicity and that ROR1 is functionally crucial for tumor growth, relapse, and metastasis. We also demonstrated that ROR1 induces activation of the E2F transcriptional network, which accelerates tumor proliferation. Epigenomic analyses identified the enhancer that supports high expression of *ROR1* through the YAP/BRD4 axis. In addition, a BET inhibitor downregulated *ROR1* and suppressed the proliferation of PDAC organoids. Our current findings indicate that intratumor ROR1^high^ cells are tumor-initiating cells in PDAC, emphasizing the implications of targeting ROR1 in PDAC therapy.

## Results

### Single-cell RNA sequencing reveals intratumor heterogeneity in a pancreatic cancer xenograft

To identify cellular diversity in PDAC, we first performed single-cell RNA sequencing (scRNA-seq) in xenograft model generated from the S2-VP10 PDAC cell line using the 10x Genomics platform (Figure 1A and Supplemental Figure 1). A total of 993 human PDAC cells were carried forward for downstream analysis after filtering low-quality and mitochondria-enriched cells (see Methods for details). We then performed clustering analysis and visualized the results using uniform manifold approximation and projection (UMAP) implemented in the Seurat package (Stuart *et al*, 2019). This analysis identified six major clusters with distinct gene expression profiles (Figures 1B–1E, and Supplemental Figure 2A and 2B): cluster 1 (cycling_G1 cells, 27.1%); cluster 2 (cycling_S cells, 17.0%); cluster 3 (cycling_G2M cells, 7.0%); cluster 4 (slow cycling cells, 26.7%); cluster 5 (autophagy cells, 8.6%); and cluster 6 (partial EMT cells, 13.6%). Cluster 4 had higher expression levels of HIF-1-regulated genes such as *NDRG1* and *VEGFA* (Ellen *et al*, 2008) (Supplemental Figure 2B), suggesting that these cells were located in hypoxic areas away from blood vessels in the tumor. In cluster 6, EMT-related markers (*ZEB1*, *MERB3*, and *VIM*) were highly expressed, while the expression of epithelial cell adhesion markers (*CDH1*, *EPCAM*, and *OCLN*) was reduced (Figure 1F). However, classical EMT-activating transcription factors (EMT TFs), such as *ZEB2*, *SNAI1*, *SNAI2*, and *TWIST1*, were barely detectable (Supplemental Figure 2C). In addition, this cluster retained the expression of epithelial origin markers, such as *KRT8* and *KRT18* (Dominguez *et al*, 2020) (Supplemental Figure 2B). Puram *et al*. (Puram *et al*, 2017) defined partial EMT as a status characterized by the expression of EMT-associated extracellular matrix genes without classical EMT TFs in head and neck cancer. Furthermore, Dongre and Weinberg (Dongre & Weinberg, 2019) recently re-defined the partial EMT state as a hybrid of epithelial and mesenchymal phenotypes. In our study, the transcriptional profile of cluster 6 cells closely resembled that of a previously reported partial EMT malignancy.

**Figure 1.**
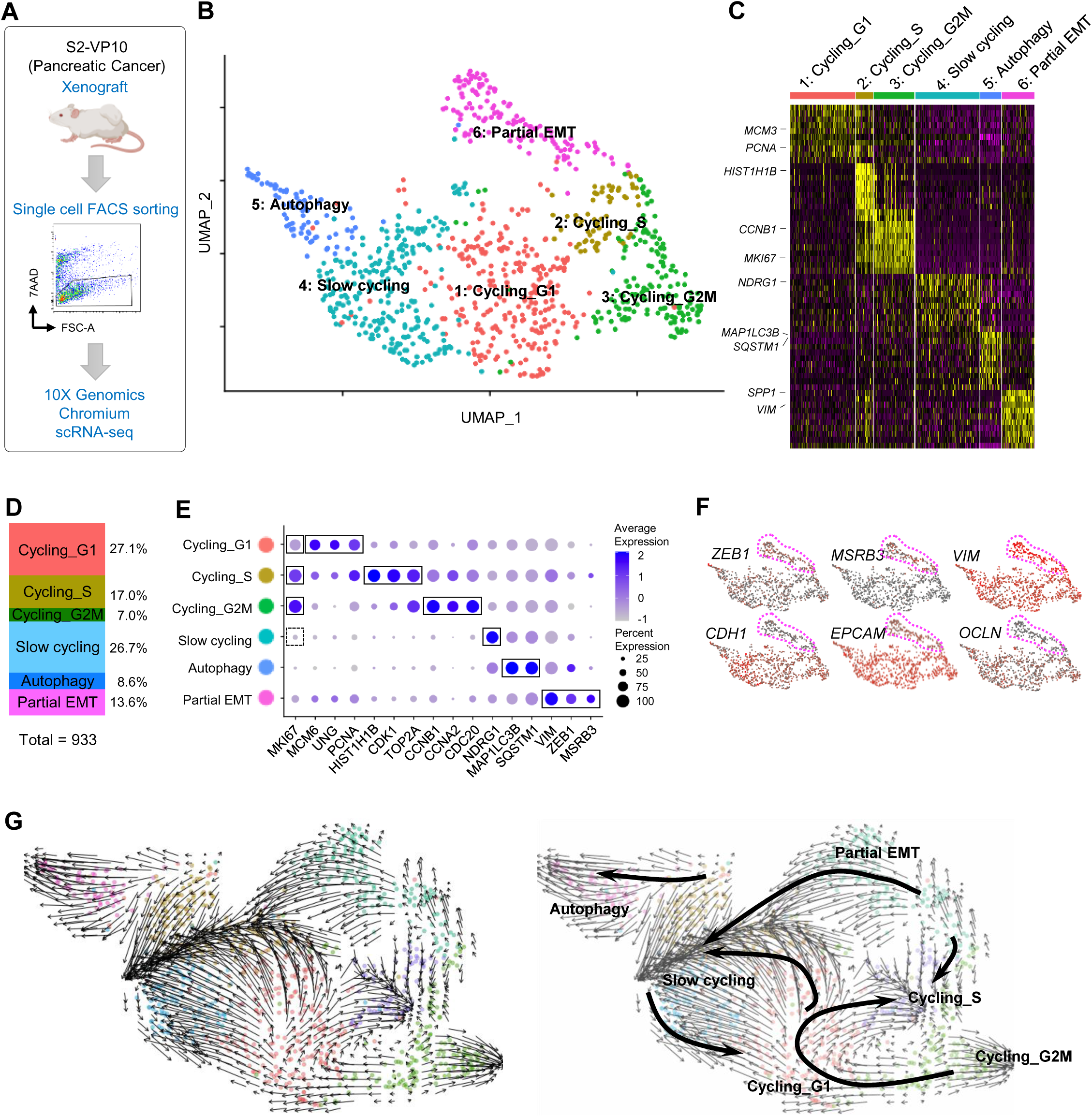
Single-cell transcriptomic analysis in pancreatic tumor xenografts. **(A)** Schematic workflow for single-cell RNA sequencing analysis using the S2-VP10 pancreatic tumor xenograft. **(B)** UMAP plot of scRNA-seq data. Six clusters of cell populations are indicated by different colors. **(C)** Heatmap showing the expression of cluster-specific genes. Representative marker genes for each cluster are indicated. **(D)** Relative proportions of each cluster. **(E)** Dot plot showing the expression levels and frequencies of marker genes in six subpopulations. The solid (high expression) and dotted (low expression) lines indicate representative markers for each cluster. **(F)** UMAP plot showing the expression of EMT markers. The magenta dotted lines indicate cluster 6 (partial EMT). **(G)** RNA velocity field projected onto the UMAP plot.

To estimate the cell lineages and potential cells of origin in the heterogeneous xenograft, we next performed RNA velocity analysis using velocyto (La Manno *et al*, 2018), a tool for predicting the future state of individual cells based on a balance between unspliced and spliced mRNAs. We found two distinct velocity flows originating from the partial EMT cluster: (1) to the cycling state, including cycling G1, cycling S, and cycling G2M, and (2) to the slow-cycling state (Figure 1G). These data suggest that the partial EMT population serves as the source of heterogeneous subpopulations in the xenograft and thus contains tumor-initiating cells.

### ROR1 marks the partial EMT population

To isolate potential tumor-initiating cells and validate their properties, we investigated the specific cell surface markers of partial EMT cells. We focused on receptor tyrosine kinases (RTKs) because aberrant activation of RTKs plays a critical role in the development and progression of cancer (Lemmon & Schlessinger, 2010). From 56 RTK genes, seven candidate genes were selected with the most enriched expression in the partial EMT cluster: *EPHA4*, *EPHA7*, *ERBB4*, *FGFR1*, *JAK3*, *LYN*, and *ROR1* (Figure 2A, Supplemental Figures 3A and 3B, and Supplemental Table 1). ROR1 is reported as an oncofetal antigen and is widely expressed in multiple human cancers (Zhang *et al*, 2012b). In addition, high expression of *ROR1* is associated with shorter metastasis-free survival in breast cancer (Cui *et al*, 2013). Therefore, we focused on ROR1 as a marker for isolating partial EMT cells (Figure 2B).

**Figure 2.**
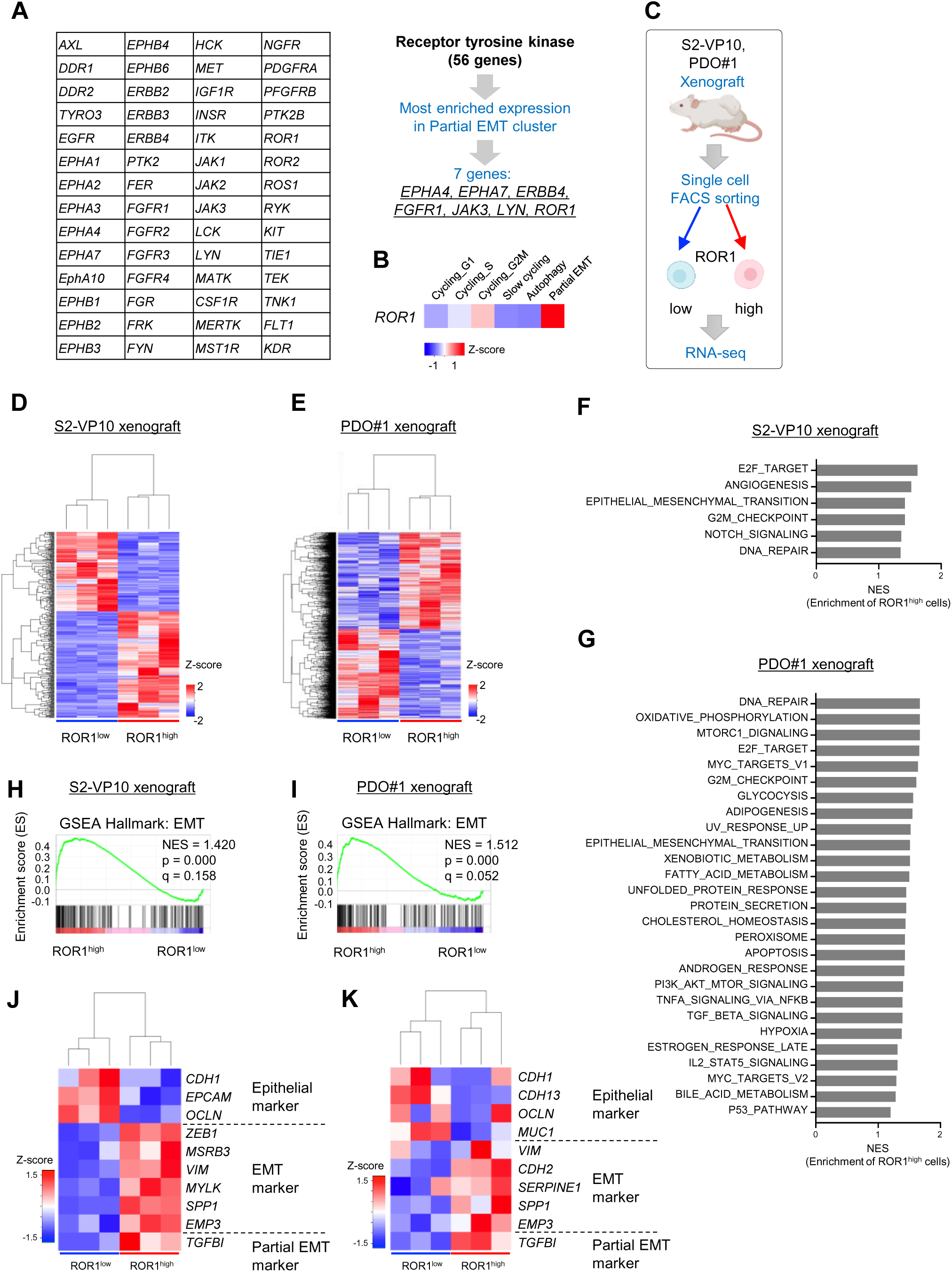
ROR1 is a surface marker for partial EMT cells in PDAC. **(A)** Strategy to identify surface markers for partial EMT cells in the S2-VP10 xenograft. **(B)** Heatmap visualizing the expression of *ROR1*. **(C)** Outline of the experimental strategy for transcriptome analysis of cells isolated from the S2-VP10 xenograft and PDO#1 xenograft (n = 3). **(D), (E)** Heatmap of DEGs in ROR1^high^ and ROR1^low^ cells in the S2-VP10 xenograft (D) or PDO#1 xenograft (E). **(F), (G)** Gene Set Enrichment Analysis (GSEA) comparing ROR1^high^ with ROR1^low^ cells (FDR < 0.25) in the S2- VP10 xenograft (F) or PDO#1 xenograft (G). NES, normalized enrichment score. See also Tables S3 and S4. **(H), (I)** GSEA plot showing significant upregulation of the EMT-related gene set in ROR1^high^ cells in the S2-VP10 xenograft (H) or PDO#1 xenograft (I). **(J), (K)** Heatmap showing epithelial, EMT, and partial EMT marker genes that are differentially expressed in ROR1^high^ versus ROR1^low^ cells in S2-VP10 xenografts (J) or PDO#1 xenografts (K).

To confirm whether ROR1 is a reliable marker of partial EMT, we isolated ROR1^high^ and ROR1^low^ cells by FACS from xenografts derived from S2-VP10 cells or patient-derived organoid (PDO) #1, and then performed RNA sequencing (Figures 2C– 2E). Gene set enrichment analysis (GSEA) (Subramanian *et al*, 2005) revealed that intratumor ROR1^high^ cells showed upregulated expression of the EMT pathway genes compared with ROR1^low^ cells (Figures 2F–2I, and Supplemental Tables 3 and 4), consistent with the scRNA-seq data. ROR1^high^ cells had higher expression of EMT-related genes but lower expression of epithelial markers than ROR1^low^ cells (Figures 2J and 2K). The partial EMT marker *TGFBI* (Puram *et al*, 2017) was also highly expressed in ROR1^high^ cells (Figures 2J and 2K). ROR1^high^ cells showed no significant upregulation of classical EMT TFs, such as *ZEB2*, *SNAI1/2*, and *TWIST1/2*, compared with ROR1^low^ cells (Supplemental Figure 3C). These results confirmed that ROR1 serves as a marker for partial EMT cells. Known CSC markers, such as *CD44*, *PROM1* (encoding CD133), and *DCLK1*, did not show a distinctive expression pattern in our scRNA-seq data (Supplemental Figure 3D).

### ROR1^high^ cells exhibit a high tumor-initiating capacity

We investigated the distribution of ROR1^high^ cells in PDAC tissue. Immunohistochemical staining showed heterogeneous expression of ROR1 in patient samples and patient-derived xenografts (PDXs) (Supplemental Figures 4A and 4B). Similarly, ROR1^high^ cells were heterogeneously present in S2-VP10 and S2-013 xenografts (Supplemental Figure 4C). Part of the tumor tissue showed a micropapillary pattern, a factor indicating poor prognosis (Reid *et al*, 2011), and was positive for ROR1 and pan-cytokeratin (an epithelial malignancy marker) (Supplemental Figure 4D).

**Figure 4.**
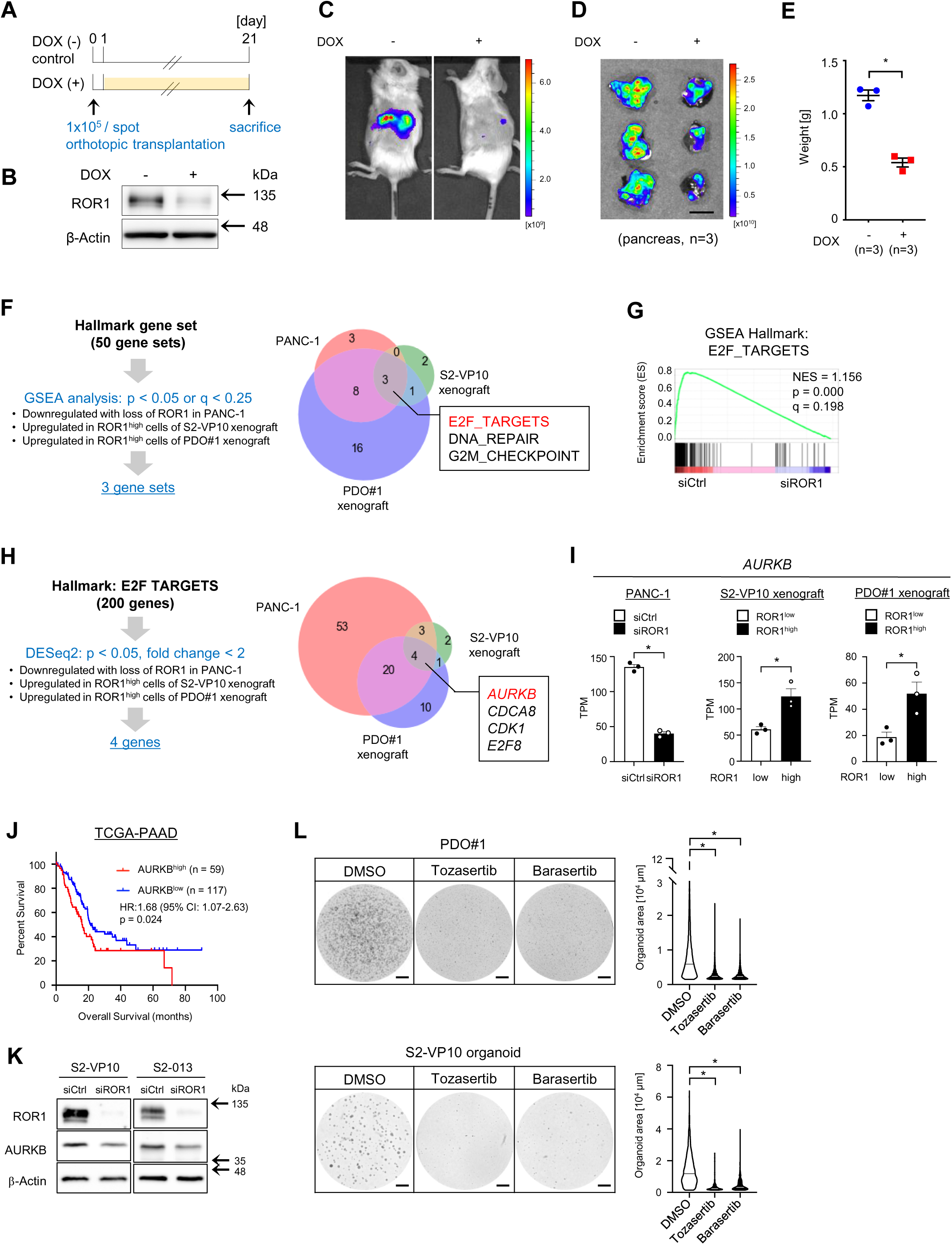
ROR1 controls AURKB via E2F activation to promote tumor proliferation. **(A)** Transplantation of mCherry-labeled S2-VP10 cells expressing doxycycline (DOX)-inducible shRNA against *ROR1* (S2-VP10-mCherry-ishROR1). **(B)** Western blot analysis of ROR1 expression in S2-VP10-mCherry-ishROR1 cells. β-Actin was used as a loading control. **(C), (D)** *In vivo* (C) and *ex vivo* (D) fluorescence imaging of S2-VP10-mCherry-ishROR1 tumors using IVIS (n = 3). **(E)** Weight of tumors derived from S2-VP10-mCherry-ishROR1 cells (n = 3). **(F)** Venn diagram showing ROR1-mediated biological states or processes. Hallmark gene sets are referenced in MSigDB. **(G)** GSEA plot showing significant downregulation of E2F activity in PANC-1 cells transfected with ROR1 siRNA. **(H)** Venn diagram to identify potential targets of ROR1 in E2F target genes. **(I)** Expression level of *AURKB* in PANC-1 cells transfected with control or ROR1 siRNA and in ROR1^high^ or ROR1^low^ cells from S2-VP10 and PDO#1 xenografts (n = 3). **(J)** Kaplan–Meier survival curves of the patients based on *AURKB* expression levels in PDAC patents form the TCGA database. **(K)** Western blot analysis of ROR1, AURKB, and β-actin in S2-VP10 and S2-013 cells transfected with control or ROR1 siRNA. **(L)** Treatment of PDO#1 and S2-VP10 organoids with DMSO (vehicle) or Aurora kinase inhibitors (r = 3). Tozasertib, pan-Aurora kinase inhibitor; Barasertib, selective Aurora B kinase inhibitor. Representative images of organoids are shown. The area of organoids is shown in the violin plot. Black or white solid lines indicate the median value for each violin. Scale bars, 1 cm (D), 1 mm (L). Data are presented as mean ± s.e.m., two-sided *t*-test. *p < 0.05.

In PDAC patients, high expression of *ROR1* was significantly associated with poor clinical outcomes in the TCGA-PAAD dataset from The Cancer Genome Atlas (TCGA) (Figure 3A), suggesting a potential role of ROR1 in PDAC progression. To investigate the tumorigenic capacity of ROR1^high^ cells, we sorted single cells from xenografts based on their ROR1 expression and examined them in two assays (Figure 3B, and Supplemental Figure 5A). (1) In Matrigel-based cultures *in vitro*, ROR1^high^ cells efficiently formed organoids or colonies compared with ROR1^low^ cells (Figure 3C and Supplemental Figure 5B). (2) An *in vivo* tumor-initiating assay showed that ROR1^high^ cells from the PDO#1 xenograft generated tumors with a higher incidence (6/6) than ROR1^low^ cells (2/6) when 500 cells were subcutaneously transplanted into *Rag2^−/−^/Jak3^−/−^*(BRJ) immunodeficient mice (Figures 3D and 3E). Similarly, ROR1^high^ cells from the S2-VP10 xenograft exhibited higher tumorigenicity than ROR1^low^ cells (Figure 3F). In addition, the xenograft derived from ROR1^high^ cells in the PDO#1 xenograft histologically recapitulated the original xenografts in terms of hierarchal morphology and cellular differentiation, such as mucus-secreting cells (Figure 3G). Thus, these data demonstrated that intratumor ROR1^high^ cells have the ability to initiate tumors and to produce differentiated progeny cells.

**Figure 3.**
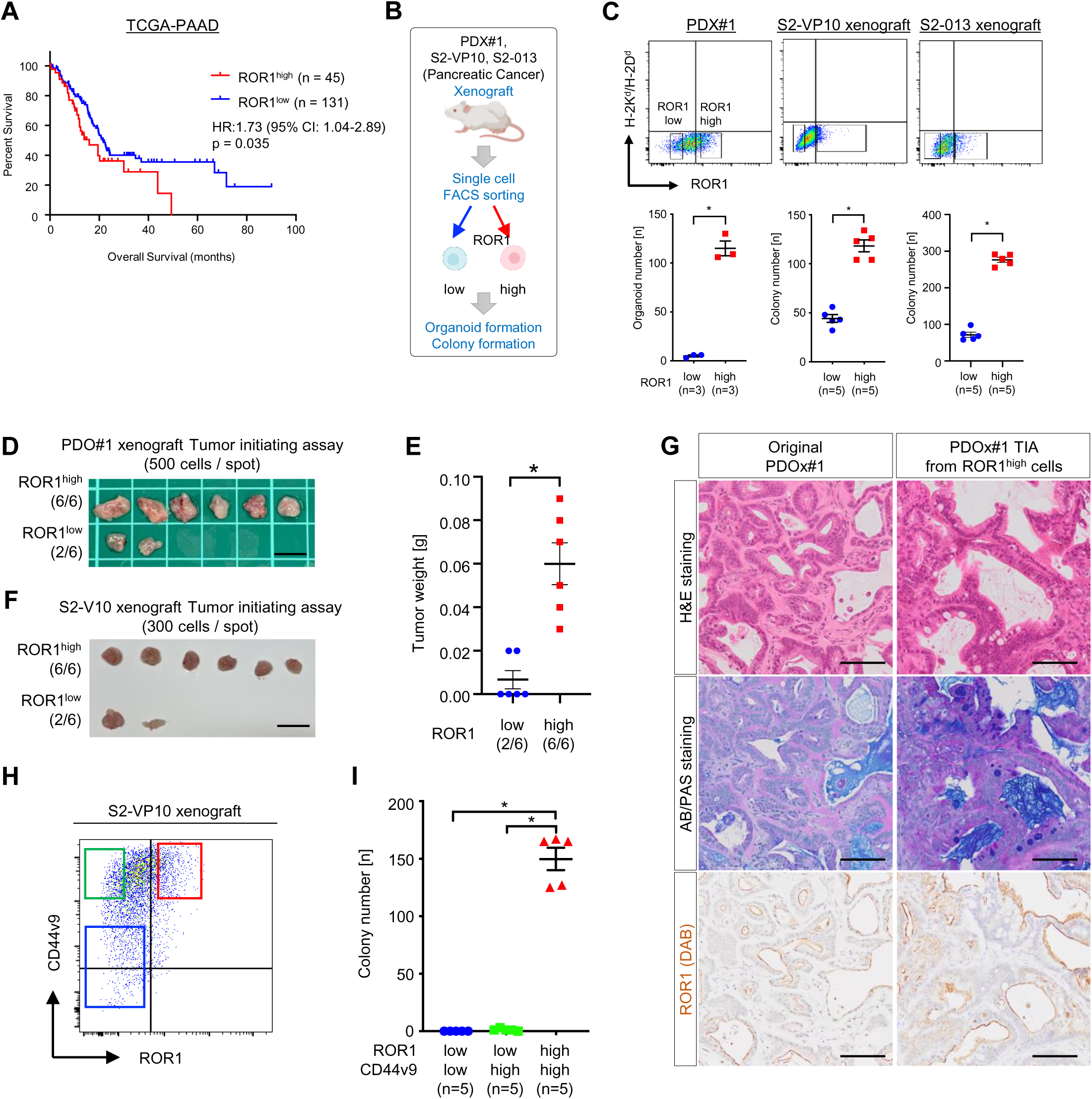
ROR1^high^ cells in PDAC have a high tumor-initiating capacity. **(A)** Kaplan–Meier survival curves of the patients based on *ROR1* expression levels in PDAC patents form the TCGA database. **(B)** Experimental strategy for the functional analyses of ROR1^high^ and ROR1^low^ cells by organoid/colony formation and tumor initiating assays. **(C)** FACS gating for sorting ROR1^high^ and ROR1^low^ cells and the number of organoids or colonies (n = 3 or n = 5). Mouse cells expressing H-2K^d^/H-2D^d^ were eliminated. **(D), (E)** Tumor initiating assay using ROR1^high^ and ROR1^low^ cells from the PDO#1 xenograft (n = 6). Images of tumors (D) and tumor weights (E) are shown. **(F)** Tumor initiating assay using ROR1^high^ and ROR1^low^ cells from the S2-VP10 xenograft (n = 6). **(G)** Representative images of tissue stained with H&E and AB/PAS, and for ROR1 in the PDO#1 xenograft and of tumor derived from ROR1^high^ cells in the PDO#1 xenograft. **(H)** FACS sorting of ROR1- and CD44v9-stained cells from the S2-VP10 xenograft. Red gating: ROR1^high^/CD44v9^high^, green: ROR1^low^/CD44v9^high^, blue: ROR1^low^/CD44v9^low^. **(I)** Colony formation assay of ROR1^high^/CD44v9^high^, ROR1^low^/CD44v9^high^, and ROR1^low^/CD44v9^low^ cells (n = 5). Scale bars, 1 cm (D), (F), and 100 µm (G) Graphs are presented as mean ± s.e.m., two-sided *t*-test. *p < 0.05.

Since CD44 is reported as a marker of tumor-initiating cells in PDAC (Li *et al*, 2007), we then investigated the relationship between ROR1 and CD44. In the S2-VP10 xenograft, FACS analysis revealed that a part of CD44^high^ cells co-expressed ROR1 (Figure 3H), and only this ROR1^high^/CD44v9^high^ population exhibited a colony-forming capability (Figure 3I and Supplemental Figure 5D). These data indicate that high ROR1 expression clearly marks tumor-initiating cells.

### ROR1 and its downstream target, AURKB, are essential for tumor growth

To investigate whether ROR1 is functionally involved in tumor growth, we generated xenografts using mCherry-labeled S2-VP10 cells expressing doxycycline (DOX)-inducible shRNA against *ROR1* (S2-VP10-mCherry-ishROR1) (Figure 4A and 4B). *ROR1*-knockdown (KD) dramatically suppressed tumor growth in mice (Figures 4C–4E), suggesting that ROR1 is not only a marker for tumor-initiating cells but is also a functional player in PDAC development.

To explore downstream effectors of ROR1 activity, we next performed an integrative analysis of three transcriptomic datasets: 1) PANC-1 cells (ROR1-expressing pancreatic adenocarcinoma cell line) transfected with control vs. ROR1 siRNA, 2) ROR1^high^ cells vs. ROR1^low^ cells in S2-VP10 xenografts, and 3) ROR1^high^ cells vs. ROR1^low^ cells in PDO#1 xenografts (Figure 4F). This analysis revealed that the E2F transcriptional network is commonly activated in ROR1-enriched samples, such as control-KD and ROR1^high^ cells, but downregulated in *ROR1*-KD and ROR1^low^ cells (Figures 4F and 4G, and Supplemental Tables 3–5). We identified four E2F target genes (*AURKB*, *CDCA8*, *CDK1*, and *E2F8*) commonly upregulated in the three ROR1-enriched datasets (Figures 4H and 4I). Aurora kinase B (AURKB) plays an important role in mitotic chromosome condensation (Lens *et al*, 2010) and has attracted considerable interest as a potential therapeutic target because of its overexpression in several cancer tissues, such as non-small cell lung carcinoma, glioblastoma, and ovarian cancer (Vischioni *et al*, 2006; Zeng *et al*, 2007; Chen *et al*, 2009). In PDAC patients, high expression of *AURKB* was significantly correlated with poor clinical outcomes (Figure 4J), suggesting a key role of the ROR1/AURKB axis in PDAC progression. We also confirmed the reduction of AURKB protein in ROR1-depleted S2-VP10 and S2-013 cells (Figure 4K). We next examined whether the kinase activity of AURKB is required for PDAC growth using PDO#1 and S2-VP10 organoids. Both tozasertib (pan-Aurora kinase inhibitor) and barasertib (Aurora kinase B selective inhibitor) markedly suppressed the growth of organoids (Figure 4L). These results indicate that AURKB is a critical downstream target of ROR1 in promoting PDAC cell proliferation.

### ROR1 depletion prevents relapse after chemotherapy

Tumor-initiating cells survive cytotoxic exposure through reversible mechanisms, leading to relapse (Boumahdi & de Sauvage, 2020). We investigated the responses of ROR1^high^ cells to drug treatment in our *in vivo* experimental model. In both the PDO#1 xenograft and S2-VP10 xenograft, gemcitabine treatment led to an increase in ROR1^high^ cells (Figures 5A and 5B), suggesting a functional involvement of ROR1^high^ cells in relapse after chemotherapy. To test this possibility, we evaluated the effect of *ROR1*-KD on tumor relapse using S2-VP10-mCherry-ishROR1 cells. After the administration of gemcitabine, tumor growth temporarily paused but resumed approximately two weeks after therapy (Figure 5C, green). In contrast, the combination of *ROR1*-KD (with Dox) and gemcitabine treatment significantly prevented relapse (Figure 5C, magenta). These results indicate that ROR1^high^ cells are a gemcitabine-resistant population and that ROR1 supports recurrence after chemotherapy.

**Figure 5.**
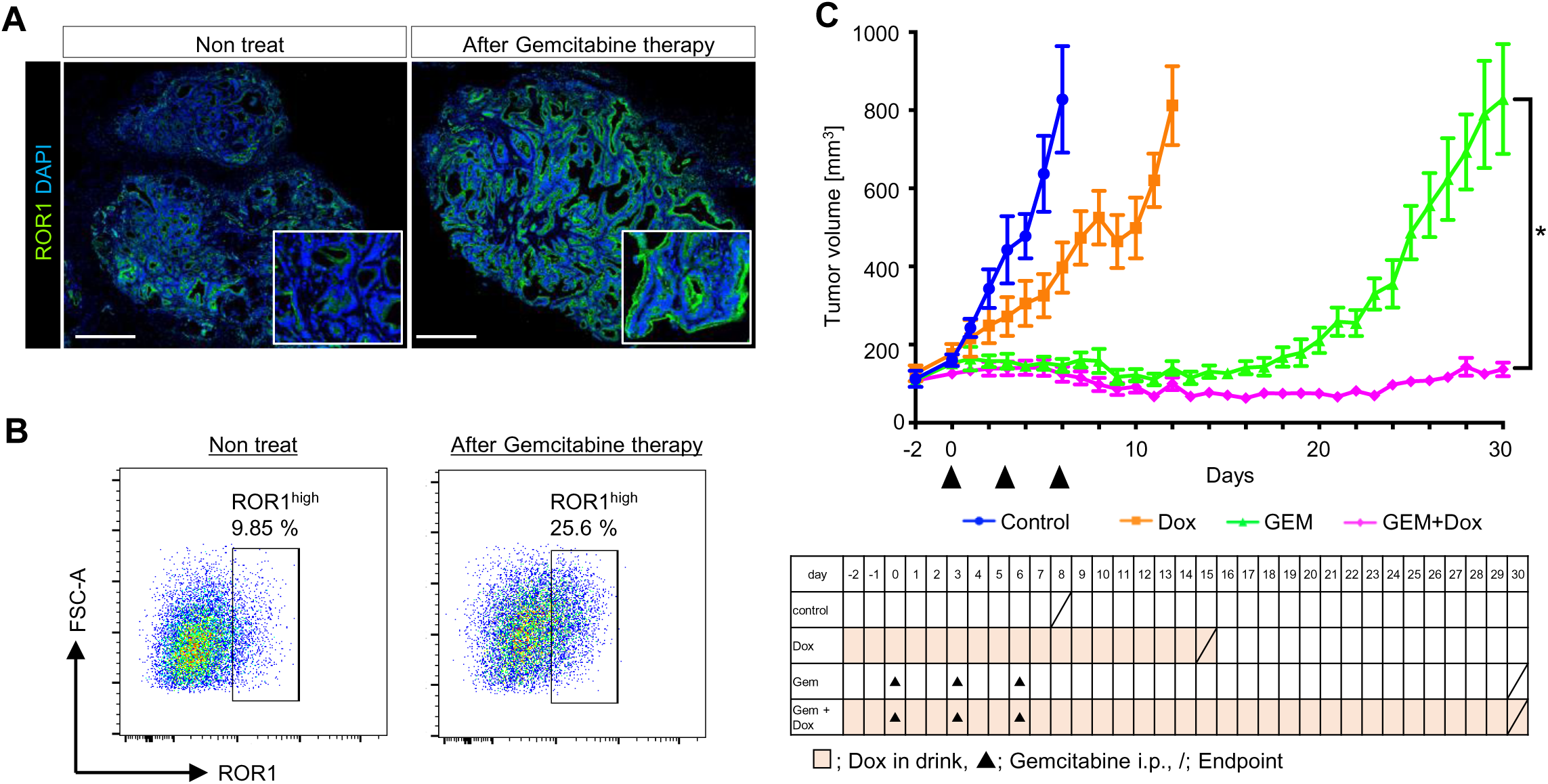
The combination of gemcitabine and *ROR1*-knockdown prevents the tumor relapse. **(A)** Representative immunofluorescence images of ROR1 in the PDO#1 xenograft with or without gemcitabine treatment. **(B)** FACS plots of ROR1 expression in cells from the S2-VP10 xenograft with or without the administration of gemcitabine. **(C)** Tumor growth curve of the four groups (control, Dox only, gemcitabine only, and Dox + gemcitabine) (n = 8). The endpoint for each group is shown by a slash. Scale bars, 500 µm (A). Data are presented as mean ± s.e.m., two-sided *t*-test. *p < 0.05.

### Inhibition of ROR1 suppresses metastasis

Tumor-initiating cells are crucial for the initiation and maintenance of metastasis (De Sousa E Melo *et al*, 2017). To investigate whether ROR1 contributes to metastasis, we prepared an orthotopically grafted mouse model in which S2-VP10-mCherry cells or S2-013 cells were transplanted into the pancreas. These cells formed primary tumors and metastases in the lung and mesenteric lymph nodes. Immunohistochemical staining revealed higher expression of ROR1 in metastatic lesions than in primary tumors (Figures 6A–6C, and Supplemental Figures 7A and 7B). In addition, ROR1^high^ metastatic foci had a higher frequency of Ki-67-positive cells than primary lesions (Figure 6D and Supplemental Figure 7C). To test whether ROR1 is required for the formation of metastatic lesions, we next examined the effect of *ROR1*-KD on metastatic potential using S2-VP10-mCherry-ishROR1 cells (Figure 6E). In the control group, multiple metastases were observed in the lung and mesenteric lymph nodes, whereas in the group treated with Dox from 10 days after transplantation, the number of metastatic foci was greatly reduced (Figures 6F and 6G). These data indicate that ROR1 is critical for inducing metastasis formation through tumor proliferation.

**Figure 6.**
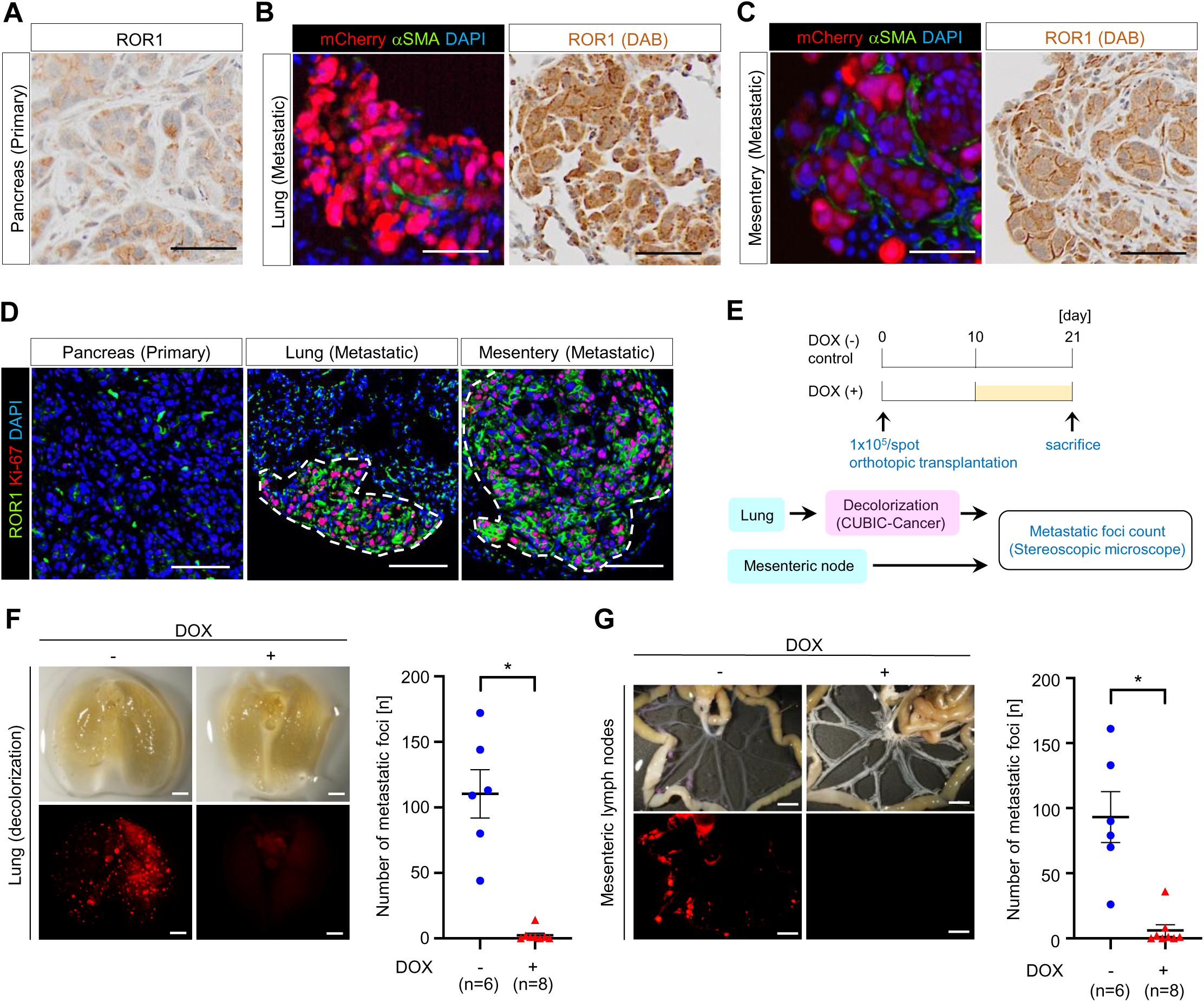
ROR1 promotes metastasis. **(A)–(C)** Representative images of ROR1, mCherry, and aSMA staining in primary tumor (pancreas) (A) and metastatic lesions in lung (B) and in mesentery lymph nodes (C) after orthotopic injection of S2-VP10-mCherry cells into the pancreas of BRJ mice. **(D)** Representative images of co-staining for ROR1 and Ki-67 in primary tumor and metastatic lesions. The dotted lines indicate the metastatic lesions. **(E)** Schematic diagram for evaluating the metastatic activity of *ROR1*-knockdown cells. **(F), (G)** Representative images and quantification of metastatic foci in lung (F) and mesentric lymph nodes (G) after orthotopic injection of S2-VP10-mCherry-ishROR1 cells into the pancreas of mice. (n = 6 or n = 8). Scale bars, 100 µm (A)–(D), 2 mm (F) and 5 mm (G). Data are presented as mean ± s.e.m., two-sided *t*-test. *p < 0.05.

### Identification of an enhancer region regulating ROR1 gene expression

As described above, we found that intratumor ROR1^high^ cells in PDAC displayed various features of tumor-initiating cells. Importantly, ROR1 functionally enhances PDAC progression, such as tumor growth, relapse, and metastasis. Therefore, we considered whether the expression of ROR1 could be druggably controlled and attempted to elucidate the epigenomic mechanisms governing *ROR1* gene expression.

We first analyzed histone modifications in cultured S2-VP10 cells that show high ROR1 expression using a CUT&RUN assay (Skene & Henikoff, 2017) and chromatin organization using an assay for transposase-accessible chromatin by sequencing (ATAC-seq). Tri-methyl histone H3 lysine 4 (H3K4me3) was enriched at the *ROR1* promoter, consistent with its actively transcribed state (Figure 7A). We also identified a putative enhancer region that opened chromatin and co-marked mono-methyl H3K4 (H3K4me1) and acetylated H3K27 (H3K27ac) at 110 kb upstream of the *ROR1* transcription start site (Figure 7A). This H3K4me1^+^/H3K27ac^+^/open chromatin region matched the active enhancer defined by the Functional Annotation of the Mammalian Genome 5 (FANTOM5) database (Figure 7A). Notably, the enrichment of H3K27ac at this region was higher in ROR1^high^ cells from the S2-VP10 xenograft than in ROR1^low^ cells (Figure 7B). Furthermore, ATAC-seq indicated higher chromatin accessibility with this region in ROR1^high^ cells than in ROR1^low^ cells (Figure 7B). To verify whether this H3K27ac^+^ region enhances transcription from the *ROR1* promoter, we cloned both regions and placed them upstream of the luciferase reporter gene (Figure 7C). The luciferase reporter assay results showed that this candidate region enhances reporter activity approximately six-fold compared with the promoter alone in S2-VP10 cells (Figure 7D). Together, these data indicate that *ROR1* transcription is strongly regulated by the newly identified enhancer, and that the difference in chromatin states at this region creates a divergence in *ROR1* expression levels among tumor cell subpopulations.

**Figure 7.**
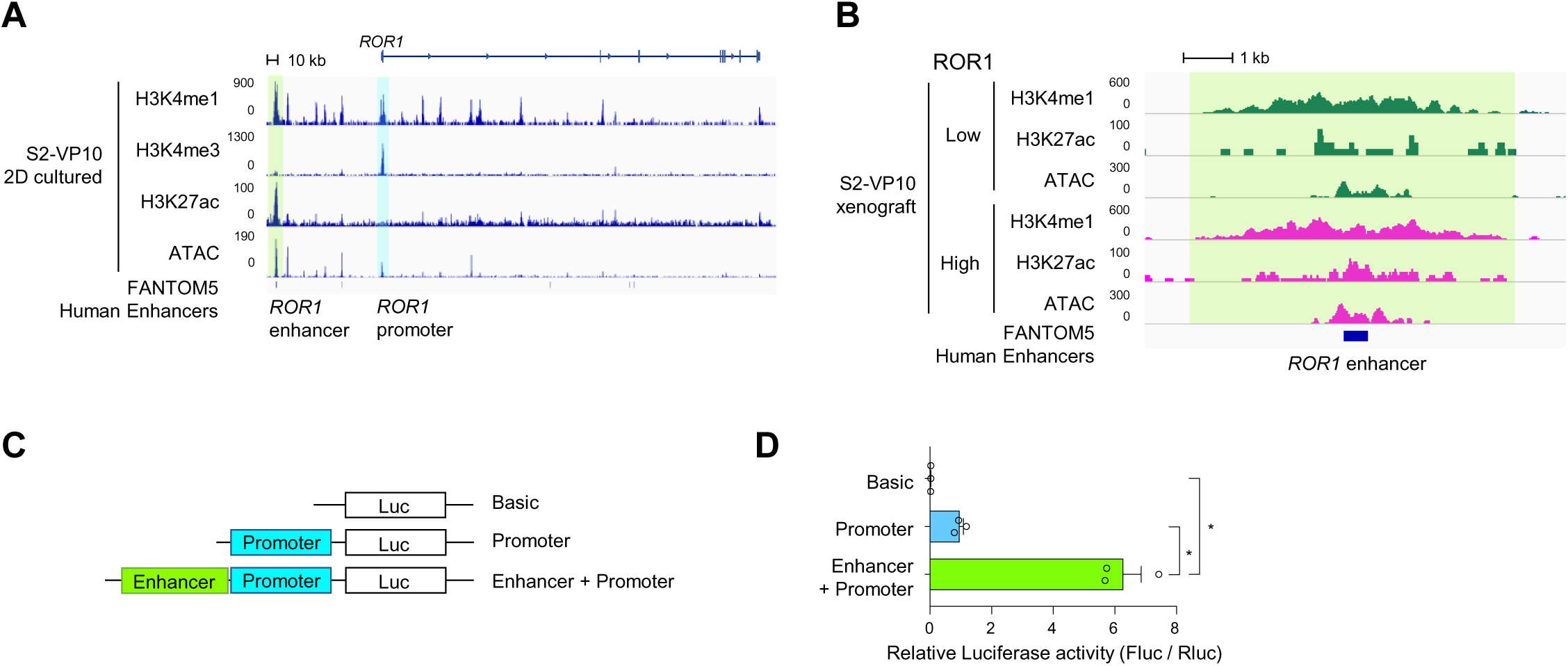
Epigenomic analyses identify the *ROR1* gene enhancer in PDAC. **(A), (B)** H3K4me1, K3K4me3, H3K27ac, and open chromatin profiles around the *ROR1* gene by CUT&RUN and ATAC-sequencing in cultured S2-VP10 cells (A) or in ROR1^high^ and ROR1^low^ cells from S2-VP10 xenografts (B). The *ROR1* promoter (blue-shaded boxes) and enhancer elements (green-shaded box) of *ROR1* are indicated. **(C)** Schematic description of vectors used for the luciferase reporter assay to examine the regions regulating the expression of the *ROR1* gene. **(D)** Relative luciferase activity in S2-VP10 cells (n = 3). Data are presented as mean ± s.e.m., two-sided *t*-test. *p < 0.05.

### ROR1 is a direct target of YAP/BRD4

We next explored the underlying mechanism for the activation of ROR1 gene expression. To identify transcription factors (TFs) that regulate the *ROR1* enhancer, we analyzed the genome-wide distribution of H3K4me1 and H3K27ac in S2-VP10 cells using a CUT&RUN assay. Using ChIP-Atlas (Oki *et al*, 2018), we compared the distribution pattern of 6887 overlapping peaks (H3K4me1 and H3K27ac) found in our study with those of TFs from available datasets (Figure 8A, Supplemental Figure 8, and Supplemental Tables 7 and 8). Of the TFs that showed a similar pattern with our peak data, we focused on Yes-associated protein (YAP). YAP and its close paralog, TAZ, are transcriptional regulators involved in CSC abilities such as tumorigenicity, chemoresistance, and metastasis in breast, esophageal, and hepatocellular cancers as well as in osteosarcoma (Bartucci *et al*, 2015; Basu-Roy *et al*, 2015; Hayashi *et al*, 2015; Song *et al*, 2014). Interestingly, both the *YAP* and *TAZ* transcript levels were significantly and positively correlated with *ROR1* transcript levels in the pancreatic adenocarcinoma dataset from the TCGA database (Figure 8B). Moreover, gene set enrichment analysis revealed that intratumor ROR1^high^ cells show upregulated expression of YAP-regulated genes compared with ROR1^low^ cells (Figure 8C). Analyses of the publicly available ChIP-seq datasets revealed that YAP binds to the enhancer region of ROR1 in human breast adenocarcinoma, lung adenocarcinoma, mesothelioma, and glioblastoma cells that express *ROR1* (Figure 8D and Supplemental Figure 9). Using ChIP-qPCR, we found direct binding of YAP at the *ROR1* enhancer in S2-VP10 cells (Figure 8E). Treatment of the cells with the YAP inhibitor verteporfin reduced the expression of *ROR1* as well as the known YAP target genes such as *CTGF* and *CYR61* (Figure 8F). Similarly, siRNA knockdown of YAP/TAZ downregulated ROR1 (Figures 8G and 8H). Together, these results indicate that YAP directly transactivates *ROR1*.

**Figure 8.**
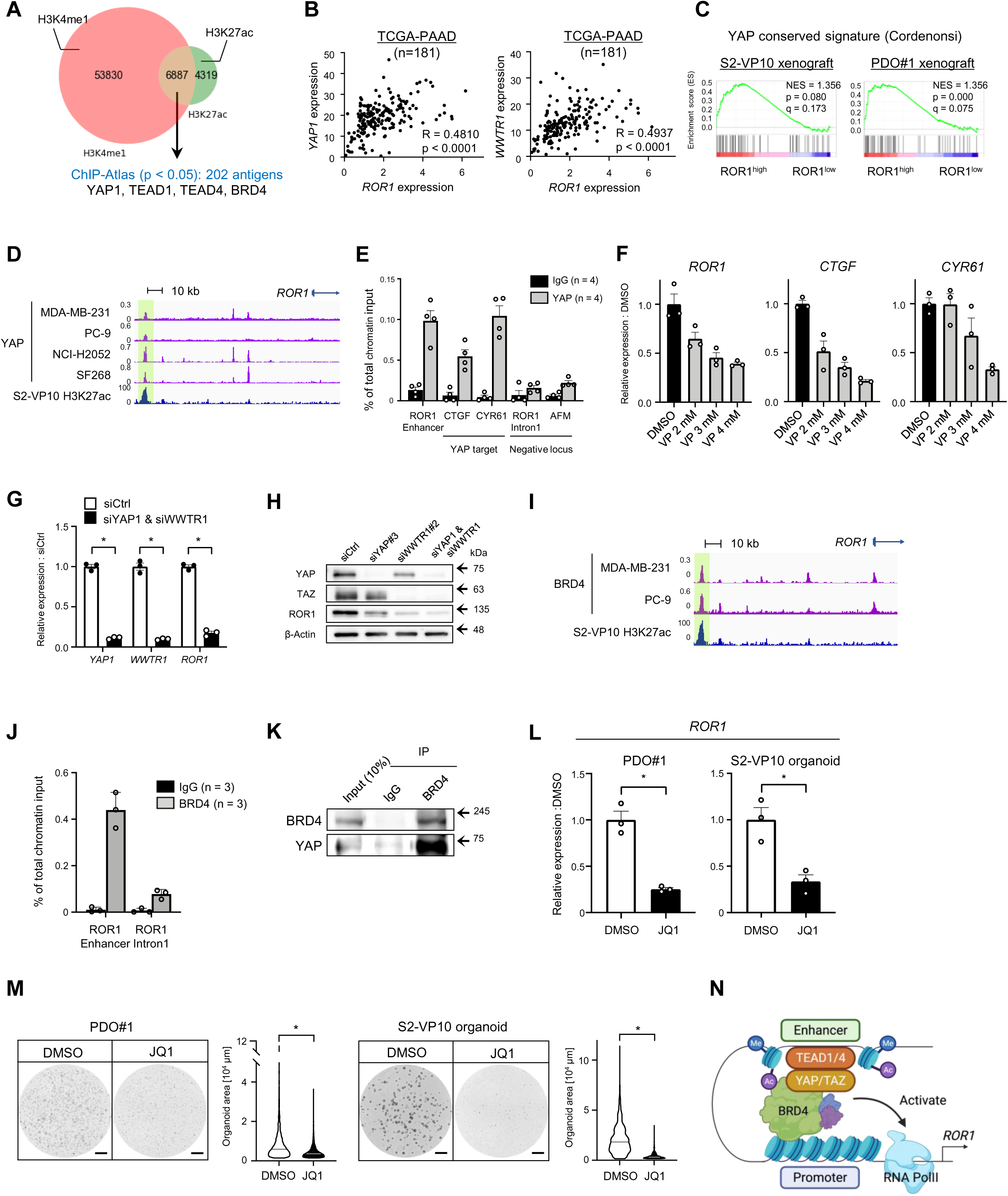
The YAP/BRD4 axis promotes ROR1 expression in PDAC. **(A)** Strategy to identify candidate antigens overlapping the H3K4me1 (salmon pink) and H3K27ac (green) peaks in S2-VP10 using ChIP-Atlas. **(B)** Pearson correlation analyses of *ROR1* expression with *YAP1* and *WWTR1* expression in PDAC patient samples (n = 181). Pearson’s correlation (R) values are indicated within each graph. **(C)** GSEA plot showing significant upregulation of the YAP conserved gene set in ROR1^high^ cells in the S2-VP10 xenograft or PDO#1 xenograft. **(D)** YAP-binding and H3K27ac profiles around the *ROR1* gene by ChIP-sequencing and CUT&RUN analyses. **(E)** YAP occupancy at the *ROR1* enhancer as determined by ChIP-qPCR. The known YAP targets, *CTGF* and *CYR61* promoters, were tested as positive controls, and *ROR1* intron 1 and *AFM* promoter were tested as negative controls (n = 4). **(F)** Relative expression of *ROR1* and YAP target genes (*CTGF* and *CYR61*) in S2-VP10 cells treated with verteporfin (VP) (n = 3). mRNA levels are normalized to that of *RPS18*. **(G)** Relative expression of *ROR1* in S2-VP10 cells transfected with YAP1 and WWTR1 siRNA (n = 3). mRNA levels are normalized to that of *RPS18*. **(H)** Western blot analysis of YAP, TAZ, ROR1, and β-actin in S2-VP10 cells transfected with control or ROR1 siRNA. **(I)** BRD4-binding and H3K27ac profiles around the *ROR1* gene detected by ChIP-sequencing and CUT&RUN analyses. **(J)** ChIP-qPCR analysis of BRD4 occupancy at the *ROR1* enhancer (n = 3). **(K)** Co-immunoprecipitation analysis to evaluate the interaction between endogenous YAP and BRD4 in S2-VP10 cells. **(L), (M)** Treatment of PDO#1 and S2-VP10 organoids with DMSO (vehicle) or a BET inhibitor (JQ1) (r = 3). Relative expression of *ROR1* (L), representative images of organoids, and organoid area (M) are shown. mRNA levels are normalized to that of *RPS18*. The area of organoids is shown in the violin plot. Black or white solid lines indicate the median value for each violin. **(N)** Schematic diagram of the regulatory mechanism of ROR1 gene expression. Created with BioRENDER.com. Scale bars, 1 mm (M). Data are presented as mean ± s.e.m., two-sided *t*-test. *p < 0.05.

ChIP-Atlas analyses also showed that bromodomain-containing protein 4 (BRD4), an acetylated histone-binding protein, has a highly similar genomic distribution with H3K4me1^+^/H3K27ac^+^ shown in our experiment (Supplemental Tables 7 and 8). BRD4 is a member of the bromodomains and extraterminal motif (BET) family (Zeng & Zhou, 2002), and BET inhibitors are rapidly being developed for clinical use because of their potent anti-tumor effects (Filippakopoulos *et al*, 2010; Doroshow *et al*, 2017). Analysis of the ChIP-seq datasets revealed the binding of BRD4 on the *ROR1* enhancer in ROR1-expressing human breast and lung adenocarcinoma cells (Figure 8I). ChIP-qPCR also confirmed the occupancy of BRD4 on the *ROR1* enhancer in S2-VP10 cells (Figure 8J). Remarkably, co-immunoprecipitation analysis revealed that BRD4 physically interacts with YAP (Figure 8K). Treatment of PDO#1 and S2-VP10 organoids with JQ1, one of the most established BET inhibitors, resulted in the downregulation of *ROR1* (Figure 8L) and the suppression of organoid growth (Figure 8M). Collectively, these data indicate that the expression of *ROR1* is regulated through the YAP/BRD4 axis (Figure 8N).

## Discussion

Understanding how tumor-initiating cells affect cancer progression and what mechanisms promote their phenomena is crucial for developing effective therapeutic strategies against malignancies. However, the tumor-initiating cells in PDAC are not fully characterized, especially at single-cell resolution. Here, we identified ROR1^high^ tumor- initiating cells in human PDAC by demonstrating their strong capability to support tumorigenicity, chemoresistance, and metastasis. Of particular note, our scRNA-seq and bulk RNA-seq analyses revealed that ROR1^high^ cells highly overlap with the partial EMT population. In skin squamous cell carcinoma (SCC), partial EMT cells characterized by the expression of CD106 and CD51 gave rise to both epithelial- and mesenchymal-like cells (Pastushenko *et al*, 2018). Another report showed that genetic ablation of the *FAT1* gene (encoding protocadherin) in skin SCC cells promoted the expression of both epithelial- and mesenchymal markers (but not CD106 and CD51), and potentiated the tumor-initiating and metastatic capacities of the cells (Pastushenko *et al*, 2021). These previous findings indicate that the partial EMT state is associated with high cellular plasticity, and that different types of partial EMT populations exist in a tumor. In the current study, we identified ROR1 as a cell surface marker for the partial EMT population with a high tumor-initiating potential in PDAC. In addition, ROR1^high^ cells enriched tumor-initiating cells from the well-known marker CD44v9^+^ CSCs. Prior studies have shown that ROR1 contributes to tumor cell survival, proliferation, migration, drug resistance, and tumorigenicity in breast and ovarian cancers (Cui *et al*, 2013; Zhang *et al*, 2014, 2019). Here, we found that ROR1 regulates AURKB levels to enhance tumor proliferation. We also demonstrated that silencing *ROR1* inhibits relapse after chemotherapy and the development of metastatic foci *in vivo*. Previous reports have shown that ROR1 induces AKT phosphorylation in breast and non-small cell lung cancers (Zhang *et al*, 2012a; Yamaguchi *et al*, 2012). Indeed, we observed a reduction of phospho-AKT levels in ROR1-depleted PDAC cells (Supplemental Figure 6). Our results clearly show that ROR1 functionally enhances the tumor-initiating potential and is thus an attractive therapeutic target in PDAC.

At a low frequency, ROR1^low^ cells also generated tumors that showed hierarchical histology mimicking the original tumor and containing ROR1^high^ cells (Supplemental Figure 5C). Interestingly, the *ROR1* enhancer employed the H3K4me1+/H3K27ac- poised chromatin state in ROR1^low^ cells (Figure 7B), suggesting that the expression of *ROR1* is flexibly regulated in PDAC. These data suggest that ROR1^low^ cells can be reversibly converted into ROR1^high^ tumor-initiating cells at a low frequency, resulting in tumor seeding. Our observation is similar to that in a previous report showing that tumor cells expressing the differentiation marker keratin 20 regain their proliferative potential and convert to LGR5^+^ CSCs in colorectal cancer (Shimokawa *et al*, 2017). Because there are different types of tumor-initiating cells (e.g., CD44^+^/CD24^+^/ESA^+^, CD133^+^, DCLK1^+^, Musashi^+^ cells), it is also possible that there are other minor tumor-initiating cells in the ROR1^low^ population.

Current therapies for PDAC are unable to ablate tumor-initiating cells effectively. Remarkably, our *in vivo* study demonstrated that treatment with gemcitabine, a conventional therapeutic agent for PDAC, enriches intratumor ROR1^high^ cells in PDAC. We also observed suppression of tumor recurrence by using a combination of gemcitabine treatment and ROR1 depletion. These findings suggest that the expansion of ROR1^high^ tumor-initiating cells after chemotherapy may be linked to efficient tumor growth during relapse. Although it remains unclear why treatment with gemcitabine enriches tumor-initiating ROR1^high^ cells, a similar phenomenon of chemotherapy-induced increase in the fraction of LGR5^+^ CSCs has been reported for colorectal and liver cancers (Osawa *et al*, 2016; Cao *et al*, 2020). Eventually, these ROR1^high^ cells may play a role in resistance against treatment and recurrence. It will be interesting to further explore the mechanisms underlying the gemcitabine-induced increase of ROR1^high^ tumor-initiating cells for anti-tumor-initiating cell therapy.

We demonstrated that the expression of *ROR1* is controlled by the enhancer with high epigenetic flexibility, leading to ROR1 heterogeneity in PDAC. Furthermore, we elucidated the mechanism of *ROR1* transactivation through the YAP/BRD4 axis. A previous study indicated that an EMT state promotes TAZ activity by inactivating Scribble, an adopter that assembles a protein complex containing MST, LATS, and TAZ, thus leading breast cancer cells to develop CSC-like traits (Cordenonsi *et al*, 2011). Our data therefore suggest that partial EMT-activated YAP/TAZ signaling maintains ROR1^high^ tumor-initiating cells. In addition, recent studies in breast cancer have revealed that ROR1 activates the Hippo-YAP pathway by phosphorylating and translocating HER3 into the nucleus (Li *et al*, 2017). Taken together, these findings suggest that ROR1 and YAP/TAZ may form a positive-feedback loop to maintain the ROR1^high^ tumor- initiating cell pool. However, the mechanism by which the transactivation of *ROR1* is restricted to particular cell populations, including tumor-initiating cells, is unclear since YAP hyperactivation is widespread in cancer tissues (Harvey *et al*, 2013; Johnson & Halder, 2014). Further studies are needed to clarify this mechanism.

ROR1 is an attractive target for cancer therapy because of its high expression in specific populations within tumors and its functional importance. As such, several therapeutic approaches are being explored against ROR1, including ROR1-targeted CAR-T therapy (Srivastava *et al*, 2019, 2021) and antibody therapy to block signals using the monoclonal anti-ROR1 antibody cirmtuzumab (Choi *et al*, 2018). Importantly, we demonstrated that Aurora kinase inhibitors and a BET inhibitor effectively suppress the growth of ROR1-mediated PDAC organoids. Our findings will greatly help in developing new therapeutic strategies for ROR1-driven PDAC. It would be fascinating to investigate in detail whether Aurora kinase inhibitors and BET inhibitors can eliminate ROR1^high^ tumor-initiating cells and suppress PDAC progression, including tumor growth, relapse, and metastasis, with minimal harmful effects on normal tissues *in vivo* studies.

The major limitation of this study is the use of xenograft models in immune-deficient mice. These models mimic clinical cancer tissue with the diversity and heterogeneity of tumors, which is helpful in investigating cancer biology, including the ability of tumor-initiating cells. However, there is a close relationship between the maintenance of tumor-initiating cells and immunosuppression in the tumor microenvironment (Bayik & Lathia, 2021). Therefore, future studies will need to use an immunocompetent mouse model to predict the therapeutic effects of targeting ROR1^high^ tumor-initiating cells accurately. Although we have demonstrated that ROR1-high expressing cells in tumors play a role in the progression of PDAC in PDX and xenograft models, whether this can also occur in cancer patients requires further investigation.

## Materials and Methods

### Mice

Male and female Balb/c;*Rag2^−/−^*/*Jak3^−/−^*(BRJ) mice were a gift from Dr. Seiji Okada (Kumamoto University) (Okada *et al*, 2011), and we used both male and female mice for this study. Female NOD.Cg-*Prkdc^scid^Il-2rg^tm1Sug^*/ShiJic (NOG) mice (7–10 weeks old) were obtained from the Central Institute for Experimental Animals (CIEA) and female C.B-17/IcrHsd-*Prkdc^scid^* (SCID) mice (7–10 weeks old) were obtained from Japan SLC. All mice were housed under specific pathogen-free conditions.

### Cell lines

Human pancreatic cancer cell lines (S2-VP10, S2-013, and PANC-1) were provided by the Cell Resource Center for Biomedical Research, Institute of Development, Aging and Cancer, Tohoku University. S2-VP10 and S2-013 cells were cultured in DMEM (low glucose) supplemented with 10% FBS at 37°C in 5% CO_2_. PANC-1 cells were cultured in RPMI-1640 supplemented with 10% FBS at 37°C in 5% CO_2_. L Wnt- 3A cells were purchased from the American Type Culture Collection. HEK293T cells were a kind gift from Dr. Toshiro Moroishi (Kumamoto University). L Wnt3A cells and HEK293T cells were cultured in DMEM (high glucose) supplemented with 10% FBS at 37°C in 5% CO_2_. All cell lines were authenticated by short tandem repeat DNA profiling and were free of mycoplasma contamination. YAP inhibitor (Verteporfin) (Cayman, 17334) was added from the next of seeding, and total RNA was extracted after 48-h incubation.

### Patient-derived xenografts (PDXs)

PDX models were established by the University of Tsukuba (Shimomura *et al*, 2018). Detailed clinical information is available in Supplemental Table 2.

### Establishment of human pancreatic cancer organoids from PDXs

Organoids were established from PDXs, based on a previous report (Seino *et al*, 2018). PDXs were washed, minced into small pieces, and digested with collagenase/dispase (Roche, 10269638001; 1 mg/mL) and DNase I (Roche, 11284932001; 1 mg/mL) in a gentleMACS Dissociator (Miltenyi Biotech) for a maximum of 60 minutes. After centrifugation at 300 × *g* for 3 minutes at room temperature (RT), the cells were treated with 1 × diluted RBC lysis buffer (BioLegend, 420301) for 30 sec and washed 3 times with ice-cold PBS containing 5% fetal bovine serum (FBS). The cell pellet was resuspended in growth-factor-reduced Matrigel (Corning, 356231) and cultured in 24-well plates (50 µL Matrigel/well) with organoid culture medium: Advanced DMEM/F-12 medium (Thermo Fisher Scientific, 12634-010) with 10 mM HEPES, 2 mM GlutaMAX-I, 1 × B27, 1 × Anti-Anti (Thermo Fisher Scientific, 15630080, 35050061, 17504044, 15240062), 10 nM Gastrin I, 1 mM N-acetylcysteine (Sigma, G9145-1MG, A9165-5G), 100 ng/mL human recombinant R-spondin-1, 100 ng/mL mouse recombinant noggin (Wako Chemicals, 181-02801, 146- 08991), 50% Wnt-3A conditioned medium from L Wnt3A cells, and 500 nM A83-01 (Tocris, 2939). For the first three days of culture, organoids were incubated in an organoid culture medium containing 10µl Y-27632 (Wako Chemicals, 036-24023). The plate was incubated in 5% CO_2_ and 20% O_2_. The medium was changed every 2 or 3 days. Organoids were passaged every 10–12 days.

### Organoid culture from cell line

300 cells of S2-VP10 cells were cultured in 50 µl Matrigel per well with S2- VP10 organoid culture medium: Advanced DMEM/F-12 medium with 10 mM HEPES, 2 mM GlutaMAX-I, 1 × B27, 1 × Anti-Anti, 10 nM Gastrin I, 1 mM N-acetylcysteine, and 100 ng/mL human recombinant R-spondin-1. S2-V10 organoids were cultured in 5% CO_2_ and 20% O_2_. 10 µL Y-27632 was added in medium for the first three days of culture.

### Organoid formation assay

10,000 ROR1^high^ or ROR1^low^ sorted cells from PDX#1, 10,000 cells of dissociated PDO#1, and 300 cells of S2-VP10 were cultured per well under the respective medium conditions. Pan-Aurora inhibitor (Tozasertib) (Selleck, S1048; 300 nM), Aurora B inhibitor (Barasertib) (Selleck, A1147; 300 nM), and BET inhibitor (JQ1) (MedChemExpress, HY13030: 100nM) were added from the day of seeding. Images of each well were captured using a BZ-X700 microscope (Keyence) on day 30 (PDX#1), day10 (PDO#1) or day 8 (S2-VP10). The organoid area was quantified using HybridCellCount software module of BZ-X Analyzer (Keyence). An area of 1,500 µm^2^ and more was identified as an organoid.

### Xenotransplantation of cell lines and organoids

The cell lines and organoids were dissociated into single cells with TrypLE Express (Thermo Fisher Scientific, 12604013). For subcutaneous transplantation, 5 × 10^5^ cells suspended in 100 µL complete medium (DMEM supplemented with 10% FBS) containing 50% Matrigel were injected into the flank of female BRJ mice or female SCID mice. For orthotopic transplantation, 1 × 10^5^ cells suspended in 50 µL complete medium containing 50% Matrigel were injected into the pancreas of anesthetized female BRJ mice.

### Single-cell isolation for scRNA-seq and flow cytometry analysis

Human PDAC xenografts were chopped and digested using collagenase/dispase (1 mg/mL) and DNase I (1 mg/mL) in a gentleMACS Dissociator for a maximum of 60 minutes. The single cell suspension was treated with 1 × diluted RBC lysis buffer and washed 3 times with ice-cold PBS containing 5% FBS. Cell suspensions were filtered using a 70 μm filter to remove debris. After centrifugation, the cells were resuspended in sorting buffer with 7-AAD (BioLegend, 420404; 1:100) to gate the viable cells. The sorting gate is shown in Supplemental Figure 1. Fluorescence-activated cell sorting (FACS) was conducted using a FACS Aria III and FACS SORP Aria (BD Biosciences).

The data were analyzed using FlowJo (version 10.6.2, BD Biosciences).

### Library preparation for scRNA-seq

The scRNA-seq libraries were generated from sorted viable single cells using Chromium Single Cell 3′ Reagent Kits v3 (10x Genomics, PN-1000092, PN-1000074, PN-120262) according to the manufacturer’s instructions. The libraries were sequenced on a Hiseq X Ten sequencer (Illumina).

### scRNA-seq data analysis

The sequencing data were mapped against the human genome (GRCh38) and quantified using the CellRanger software package (version 6.0). Raw gene counts were imported into R (version 4.0.2) and processed using the R package Seurat (version 4.0.4). Cells with <1% and >30% mitochondrial genes mapped, <5000 count RNA (UMIs), and <4000 feature RNA (expressed genes) were eliminated from the downstream analysis. After filtering, UMI counts were normalized and subjected to principal component analysis (PCA). Clustering analysis was performed using the *FindNeighbors* and *Findclusters* functions of Seurat with the parameters Findneighbors (dim = 15) and Findclusters (resolution = 0.7). Dimensional reduction was performed by UMAP. RNA velocity analysis was carried out using Velocyto.R (version 0.6) with default parameters.

### Flow cytometry

After dispersal of the xenografts into single cells, the cell pellet was suspended in sorting buffer (D-PBS with 2% FBS, 1 mM EDTA, and 25 mM HEPES). The cells were first incubated with anti-ROR1 antibody (BD Biosciences, 564464) and/or anti- CD44v9 antibody (Cosmo Bio, LKG-M003) at 2 μg/mL and incubated on ice for 30 min.

After washing, the cells were stained with goat anti-mouse IgG_2b_ labeled Alexa Fluor 647 (Thermo Fisher Scientific, A21242; 1:1000) or goat anti-rat IgG (H+L) labeled PE (Thermo Fisher Scientific, A10545; 1:200) and PE/Cy7-labeled anti-H-2K^d^/H-2D^d^ (BioLegened, 114718; 1:50) on ice for 30 min in the dark. After washing twice, the cells were resuspended in sorting buffer with 7-AAD (BioLegend) to gate the viable cells. The sorting gate is shown in Supplemental Figure 6A. Flow cytometric analysis was performed using FACSVerse (BD Biosciences), and FACS was conducted using a FACS Aria III and FACS SORP Aria (BD Biosciences).

### RNA-seq analysis

For the S2-VP10 xenograft, RNA was extracted from sorted ROR1^low^ and ROR1^high^ cells using a MagMAX-96 Total RNA Isolation Kit (Thermo Fisher Scientific, AM1830), and 1 ng of RNA was used for the reverse transcription reaction using SMART-seq HT (Takara, 634455). For the PDO xenograft, ROR1^low^ and ROR1^high^ cells were collected directly into CDS Sorting Solution by FACS sorting, then cDNA was synthesized without RNA purification using SMART-seq HT (Takara). For the PANC-1 cells, RNA was extracted using total RNA was extracted using a ReliaPrep RNA Miniprep System (Promega, Z6012), and 1 ng of RNA was used for the reverse transcription reaction using SMART-seq HT. The RNA-seq library was prepared using a Nextera XT Library Prep Kit (Illumina, FC-131-1024, FC-131-1001) and sequenced using a Hiseq X Ten sequencer (Illumina). RNA-seq reads were aligned to a human transcriptome (GRCh38) and quantified by Salmon (Patro *et al*, 2017) (version 1.3.0) with default settings. Differential expression testing was performed with DESeq2 (Love *et al*, 2014) (version 1.28.1). A p-value threshold of <0.05 was used to determine differentially expressed genes. Gene set enrichment analysis was performed using GSEA (version 4.0.3) (Broad Institute) with H hallmark gene sets in the Molecular Signatures Database. The heat maps were drawn using the ggplot2 R package.

### Real-time quantitative PCR

For cultured S2-VP10 cells, total RNA was extracted using a ReliaPrep RNA Miniprep System (Promega, Z6012), and cDNA was synthesized using a High Capacity cDNA Reverse Transcription Kit with RNase Inhibitor (Thermo Fisher Scientific, 4374967). Real-time qPCR reactions were performed on a ViiA 7 Real-Time PCR System (Thermo Fisher Scientific) using PowerUP SYBR Green Master Mix (Thermo Fisher Scientific, A25742). Samples were run in three biological replicates, and mRNA levels were normalized to *RPS18*. The primers used are listed in Supplemental Table 9.

### Histology and immunostaining

Xenografts were isolated and immediately fixed in 4% paraformaldehyde (Nakalai Tesque, 09154-56) at 4°C for 16 h. Paraffin-embedded samples were cut into 4-μm sections and used for histological analysis. Hematoxylin and eosin (H&E) staining and alcian blue and periodic acid-Schiff (AB/PAS) staining were performed using standard protocols. For immunohistochemistry (IHC), antigen retrieval was carried out in an autoclave (2100 Retriever, Aptum Biologics) using 0.01 M citrate buffer (pH 7.0).

After treating with 3% H_2_O_2_ to block endogenous peroxidase activity, sections were incubated in Tris-buffered saline/0.1% Tween-20 (TBS-T) containing 5% goat serum (Jackson Immuno Research Laboratories, 005-000-001) to block non-specific binding. The following primary antibodies were used: rabbit anti-ROR1 (Thermo Fisher Scientific, PA5-50830; 1:50), rabbit anti-pan cytokeratin (Nichirei, 422061; 1:2), rabbit anti-RFP (MBL, PM005; 1:400), and mouse anti-alpha smooth muscle actin (Thermo Fisher Scientific; 14-9760-82; 1:400). The secondary antibodies were horseradish peroxidase- conjugated goat anti-rabbit IgG antibodies (Nichirei, 424141) for HRP-IHC and visualized using a Liquid DAB+ Substrate Chromogen System (DAKO, K3468), or Alexa Fluor Plus 488- or 555-conjugated goat anti-mouse, rabbit IgG antibodies (Thermo Fisher Scientific, A32723, A32731, A32727, A32732; 1:1000) for immunofluorescence. Nuclei were counterstained with hematoxylin or DAPI (Dojindo, 340-07971). All slides were imaged using a BZ-X700 fluorescence microscope or VS120 fluorescence virtual slide microscope (Olympus).

### Colony formation assay

For 2,000 sorted cells, 20 μL of 50% Matrigel containing complete medium was added and mixed gently on ice. A 96-well plate was chilled on ice and coated with 25 μL of cell-free 50% Matrigel. After polymerization of the coated Matrigel at 37°C for 10 min, the cell mixture was overlaid at a density of 2,000 cells per well. Half the volume of the medium was changed every 2–3 days. Images of each well were captured using a BZ-X700 microscope on day 8.

### Tumor-initiating assay

Sorted ROR1^low^ and ROR1^high^ cells from xenografts were mixed with 50% Matrigel containing complete medium. The cell suspension was subcutaneously injected into the flank of female BRJ mice or female NOG mice. The resulting tumors were harvested after 56 days (PDO#1 xenograft) or 27 days (S2-VP10 xenograft) post-injection.

### Lentivirus preparation and establishment of stable cell lines

To generate lentivirus-based doxycycline inducible shRNA constructs against *ROR1*, the shRNA sequences (5′-CTCATTTAGCAGACATCGCAA-3′ (shRNA-1) and 5′-CTTTACTAGGAGACGCCAATA-3′ (shRNA-2) (Zhang *et al*, 2012b)) were inserted into EZ-Tet-pLKO-Puro vector (Addgene, 85966), respectively. The preparation of lentivirus has been described previously (Daikuzono *et al*, 2021). Cells were infected with the two lentiviral shRNAs against *ROR1* and selected with 0.5 μg/mL puromycin (Nakalai Tesque, 29455-12) for 48 h. To induce the expression of shRNA *in vitro*, 1 μg/mL doxycycline (Tokyo Chemical Industry, D4116) was added to the cells. For *in vivo* studies, mice were given drinking water supplemented with 2 mg/mL doxycycline (Tokyo Chemical Industry) and 5% sucrose (Wako Chemicals, 196-00015). Cells were labeled with mCherry using the pLV-mCherry vector (Addgene, 36084). EZ-Tet-pLKO- Puro was a gift from Cindy Miranti (Frank *et al*, 2017). pMD2.G and psPAX2 were a gift from Didier Trono. pLV-mCherry was a gift from Pantelis Tsoulfas.

### In vivo fluorescence imaging of tumor growth

*In vivo* fluorescence imaging was performed using an IVIS SPECTRUM (Caliper Life Sciences). The mCherry-labeled tumor-bearing mice were anesthetized with isoflurane and imaged. The filter settings were Ex = 570 nm, Em = 620 nm. Data were quantified with Living Imaging software (version 4.3.1).

### Western blotting

Cells were lysed with RIPA buffer (Wako Chemicals, 182-02451) containing cOmplete protease inhibitor cocktail (Roche, 05056489001) and PhosSTOP phosphatase inhibitor cocktail (Roche, 05056489001). Whole-cell lysate was loaded and separated on a 7.5% polyacrylamide gel and transferred onto a PVDF membrane (Merck Millipore, IPVH07850). Membranes were blocked with PVDF Blocking Reagent for Can Get Signal (Toyobo, NYPBR01) and then incubated with primary antibody in Can Get Signal Solution 1 (Toyobo, NKB-101). The following primary antibodies were used: mouse anti- β-actin (Sigma, A1978; 1:10,000), rabbit anti-ROR1 (Cell Signaling Technology, 16450; 1:250), rabbit anti-AKT (Cell Signaling Technology, 4691; 1:1000), rabbit anti-phospho-AKT (Ser473) (Cell Signaling Technology, 9271; 1:1000), rabbit anti-Aurora B (abcam, ab2254; 1:1000), rabbit anti-YAP (Cell Signaling Technology, 14074; 1:1000), rabbit anti-TAZ (Cell Signaling Technology, 72804; 1:1000), and rabbit anti-BRD4 (Cell Signaling Technology, 13440; 1:1000). Signals were detected by HRP-conjugated anti-rabbit or mouse antibody (Cell Signaling Technology, 7074, 7076; 1:1000) in Can Get Signal Solution 2 (Toyobo) and visualized with ECL Prime Western Blotting Detection Reagent (GE Healthcare, RPN2232). Chemiluminescent signals were detected using a LAS-3000 Imaging System (GE Healthcare).

### RNA interference

Reverse transfection of siRNA was performed using lipofectamine RNAiMAX (Thermo Fisher Scientific, 13778150) with a final siRNA concentration of 10 nM. Cells were harvested after 48-h incubation. The siRNAs were purchased from Qiagen and Dharmacon (Qiagen, negative control siRNA: 1027280, siROR1: SI00071295; Dharmacon, negative control siRNA: D-001810-01, siYAP1#3:J-012200-07-0002, siWWTR1#2: J-016083-06-0002).

### Xenograft experiments with recurrent and metastasis models

For the recurrence experiments, mice implanted with S2-VP10 cells (14 days post-implantation) or PDO#1 (18 days post-implantation) were given gemcitabine (Pfizer, 4987114700506; 120 mg/kg) intraperitoneally 3 times daily for 3 days. For flowcytometric analysis and IHC analysis, xenografts were harvested 1 day after the last administration. Tumor volume (Tv) measurements were made every day (Tv [mm^3^] = (*length* [mm] × *width* [mm]^2^)/2).

For the metastasis experiments, mice implanted with S2-VP10 cells or S2-013 cells orthotopically into the pancreas were sacrificed at 21 days post-implantation. The metastatic foci in the isolated lung and mesentery were counted using a SZX12 stereo microscope (Olympus). To observe tissue deeply, lung samples were decolorized using CUBIC-Cancer method (Kubota *et al*, 2017).

### CUT&RUN

CUT&RUN was performed using a CUT&RUN Assay Kit (Cell Signaling Technology, 86652) with a modification of the manufacturer’s protocol. Between 28,000–50,000 collected cells were washed and bound to activated concanavalin A-coated magnetic beads. After permeabilization, the bead-cell complex was incubated with antibody at RT for 2 h on a nutator, with gentle tapping every 20 min. After washing, the beads were resuspended in pAG-MNase and incubated at 4°C for 1 h, with gentle tapping every 20 min. After washing, the beads in digitonin buffer were chilled in ice water bath (0°C), then pre-cooled calcium chloride was added to activate MNase for 30 min. Enriched DNA fragments were purified by phenol-chloroform-isoamyl alcohol/ethanol extraction. The following antibodies were used: rabbit anti-mono-methylated histone H3K4 (Abcam, ab8895; 1 μg), rabbit anti-tri-methylated histone H3K4 (Millipore, 07- 473; 1 μg), rabbit anti-acetylated histone H3K27 (Cell Signaling Technology, 8173; 1:100), and rabbit isotype control IgG (BioLegend, 910805; 1 μg). The CUT&RUN library was prepared using a NEBNext Ultra II DNA Library Prep Kit for Illumina (New England Biolabs, E7645, E7710) as previously reported (Zhu *et al*, 2019). High-throughput sequencing was performed using a NextSeq 500 Sequencer with 75-bp single-end reads. Qualified reads were aligned to the human genome (GRCh37) using Bowtie2 (Langmead & Salzberg, 2012) (version 2.3.4.1). Duplicate reads were removed. The number of unique reads was normalized to spike-in yeast DNA (sacCer3). The final number of mapped reads and scaling factors are listed in Supplemental Table 6. Peak detection was performed using MACS2 (Zhang *et al*, 2008) (version 2.2.7.1) with default parameters. The overlapping peaks between H3K4me1 and H3K27ac were obtained using the bedtools intersect function (version 2.30.0). CUT&RUN data were converted to bigwig files by bamCoverage (in the deepTools package (Ramírez *et al*, 2016), version 3.5.1) using the parameters --scaleFactor <each sample> --binSize 10, then visualized using Integrative Genome Viewer (IGV). Enrichment analysis of the overlapping peaks (H3K4me1 and H3K27ac) with other TFs was performed using ChIP-Atlas with the following parameters: [Experiment type] ChIP TFs and others; [Cell type Class] Pancreas, Breast, Lung; [Threshold for Significance] 100. The aggregation plot was drawn using computeMatrix and plotHeatmap of deepTools.

### ATAC-sequencing

ATAC-sequencing was performed with modifications based on a previous report (Corces *et al*, 2017). The collected cells (2D cultured: 50,000 cells, sorted from xenografts: 15,000 cells) were suspended in 50 µL of cold cell lysis buffer (10 mM Tris-HCl pH 7.5, 10 mM NaCl, 3 mM MgCl_2_, 0.1% NP-40, 0.1% Tween-20, 0.01% digitonin in water) by pipetting three times and incubated on ice for 3 min. After lysis, 1 mL wash buffer (10 mM Tris-HCl pH 7.5, 10 mM NaCl, 3 mM MgCl_2_, 0.1% Tween-20) was added, and the reaction mixture was centrifuged at 500 × *g* for 10 min at 4°C to collect nuclei. The supernatant was removed, and the nuclei were resuspended in 50 µL transposition mixture (25 µL 2× Tagmentation buffer (Diagenode, C01019043), 2.5 µL Tagmentase loaded (Diagnode, C01070012), 0.5 µL digitonin (final concentration 0.01%), 0.5 µL Tween-20 (final concentration 0.1%), 16.5 µL PBS, and 5 µL nuclease-free water). The transposition reaction mixture was incubated for 30 min at 37°C in a block incubator. The transposition DNA fragments were purified using a MinElute PCR Purification Kit (Qiagen, 28006) and eluted in 20 µL buffer EB. Transposed DNA fragments were amplified by combining the following: 20 µL transposed DNA, 2.5 µL of 25 μM PCR Primer 1 (i5), 2.5 µL of 25 μ PCR Primer 2 (barcoded, i7), and 25 µL KAPA HiFi HS Ready Mix (Kapa Biosystems, KK2601). The following thermal cycler conditions were used: 72°C for 5 min, 98°C 30 sec, followed by 11 cycles (50,000 cells) or 12 cycles (15,000 cells) of 98°C 10 sec, 63°C 30 sec, and 72°C 1 min. The amplified library was purified using a 1.8× volume of AMPure XP beads (Beckman Coulter, A63880) and eluted in 20 µL buffer EB. To remove large-size fragments (>1000 bp), a 0.55× volume (11 µL) of SPRIselect (Beckman Coulter, B23317) was added to the purified library. Following 2-min incubation at room temperature, the reaction tube was placed in a magnetic rack for 5 min. After the liquid appeared completely clear, 31 µL of the supernatant was transferred to a new tube and 25 µL SPRIselect (final volume of 1.8× based on the initial reaction volume) was added to the sample. After 2-min incubation at room temperature, the library was washed twice with 80% ethanol and eluted in 20 µL buffer EB. Library size distributions were checked using an Agilent 2200 TapeStation (Agilent Technologies). High-throughput sequencing and data analysis were performed by the same method as CUT&RUN. ATAC-seq data were converted to bigwig files using bamCoverage with default parameters. The primers used are listed in Supplemental Table 9.

### FANTOM5 enhancer data

Human enhancers were searched using FANTOM5 Human Enhancers Selector (https://slidebase.binf.ku.dk/human_enhancers/selector) using the parameters Genes ROR1, Upstream 1,000,000 bp, and Downstream 150,000 bp.

### Luciferase reporter assay

S2-VP10 genomic DNA was extracted using a DNeasy Blood & Tissue Kit (Qiagen, 69504). The promoter and enhancer candidate regions of ROR1 were amplified by PCR using KOD -Plus- Neo (Toyobo, KOD-401) and subcloned into the pGL3 luciferase reporter vector (Promega, E1751) (Figure 6C). The primers used are listed in Supplemental Table 9. All inserts were confirmed by sequencing. For transient transfection, S2-VP10 cells were co-transfected with 0.5 μg of each luciferase reporter plasmid and 1 ng of pRL-SV40 (Promega, E2231) the day after plating (3.45 × 10^4^ cells/well in a 24-well plate) using FuGENE HD (Promega, E2312). After 48-h incubation, the cells were harvested and luciferase was measured using a Dual-Luciferase Reporter Assay System (Promega, E1980). The promoter and enhancer activity were calculated by the ratio of firefly/renilla luciferase activity.

### ROR1 and YAP/TAZ gene expression analysis (TCGA database)

Gene expression data were downloaded from TCGA (TCGA-PAAD, Retrieved on March 22, 2021). *ROR1* and *YAP1/WWTR1* expression were extracted and evaluated by Pearson correlation analysis. Correlation coefficients and p-values are reported.

### ChIP-seq data analysis

Bigwig files were retrieved from the GEO database (GSE66081, GSE131687, GSE61852, GSE116879, and GSE89128). *ROR1* gene expression data were downloaded from the CCLE database. All data were visualized using IGV.

### ChIP-qPCR

About 5 × 10^6^ S2-VP10 cells were used to detect YAP and BRD4 enrichment. Cells were crosslinked with 1% formaldehyde for 10 min at RT, then equilibrated with 0.135 M glycine. Cells were lysed in lysis buffer (5 mM PIPES [pH 8.0], 85 mM KCl, 5% NP-40). Isolated nuclei were suspended in low-salt SDS buffer (0.1% SDS, 10 mM EDTA, 50 mM Tris-HCl [pH 8.0]) and sonicated to fragment the chromatin using a Bioruptor UCD-300 (high, 30 sec on/30 sec off, 20 min, Cosmo Bio). After preclearing at 4°C for 1 h, chromatin fragments were incubated at 4°C overnight with antibodies for YAP (Cell Signaling Technology; 1:100) or BRD4 (Cell Signaling Technology; 1:100). Antibody/antigen complexes were recovered with Dynabeads M-280 sheep anti-rabbit IgG (Thermo Fisher Scientific, 11203D). Beads with DNA fragments were washed once with RIPA buffer (50 mM Tris-HCl, 150 mM NaCl, 1 mM EDTA, 1% Triton X-100, 0.1% SDS, 0.1% sodium deoxycholate), twice with high salt RIPA buffer (50 mM Tris-HCl, 500 mM NaCl, 1 mM EDTA, 1% Triton X-100, 0.1% SDS, 0.1% sodium deoxycholate), once with LiCl wash buffer (10 mM Tris-HCl, 250 mM LiCl, 1 mM EDTA, 0.5% NP-40, 0.5% sodium deoxycholate), and twice with TE buffer. The washed beads with DNA fragments were resuspended in ChIP elution buffer (0.5% SDS, 10 mM Tris-HCl, 5 mM EDTA, 300 mM NaCl) and incubated at 65°C for 6 h to reduce the crosslinks. Enriched DNA fragments were purified by phenol-chloroform-isoamyl alcohol/ethanol extraction and subjected to qPCR analysis. The primers used are listed in Supplemental Table 9.

### Co-immunoprecipitation of endogenous nuclear proteins

About 1 × 10^7^ S2-VP10 cells were used for the analysis. Proteins were crosslinked using 1 mM dithiobis (succinimidyl propionate) (DSP) (Dojindo, D629) to increase the stability of the protein-protein complexes. The cells were rinsed with ice-cold PBS and harvested using a cell scraper. The nuclei were isolated in nuclear extraction buffer (20 mM HEPES-KOH (pH 8.0), 10 mM KCl, 0.1% NP-40, 20% glycerol, freshly added protease inhibitors) at 4°C for 15 min. After centrifugation and discarding the supernatant, the nuclear pellet was lysed in hypertonic buffer (20 mM HEPES, 400 mM NaCl, 1 mM EDTA, 0.5% NP-40, freshly added protease inhibitors) and sonicated using a Bioruptor UCD-300 (high, 30 sec on/30 sec off, 10 min). The nuclear lysates were centrifuged at 18,000 × *g* for 10 min at 4°C, and the supernatant was collected and diluted in 150 mM NaCl. After preclearing for 1 h at 4°C, the lysates were incubated at 4°C for 5 h with antibodies for BRD4 (Cell Signaling Technology; 1:100). Antibody/antigen complexes were recovered using Dynabeads Protein A/G (Thermo Fisher Scientific, DB10001, DB10003) for 1 h at 4°C. The immunocomplex was washed four times with wash buffer (20 mM HEPES, 150 mM NaCl, 1 mM EDTA, 0.1% NP-40, freshly added protease inhibitors). Proteins were eluted in SDS sample buffer with 2-mercaptoethanol at 95°C for 5 min and subjected to western blot analysis.

### Survival analysis (TCGA database)

The gene expression data and clinical data were downloaded from TCGA (TCGA-PAAD, Retrieved on March 22, 2021). The association between the expression of *ROR1* and patient overall survival was examined by the Kaplan–Meier method. The log-rank (Mantel–Cox) test was used to assess statistical significance in overall survival between patients with *ROR1^high^*and *ROR1^low^* or *AURKB^high^* and *AURKB^low^*.

### Statistics

Pairwise comparisons were performed using an unpaired two-tailed Student’s *t*- test. Results are presented as the mean ± s.e.m. (standard error of the mean). A p-value less than 0.05 was considered statistically significant. Statistical analysis was performed using Prism 8 (GraphPad Software Inc.).

### Study approval

All animal experiments in this study were performed based on protocols approved by the Institutional Animal Care and Use Committee of Kumamoto University, Japan. Clinical samples were obtained from patients at University of Tsukuba Hospital with informed consent after approval by the ethical committees. All human experiments were approved by University of Tsukuba and Kumamoto University.

### Data availability

The accession number for the scRNA-seq, RNA-seq, CUT&RUN, and ATAC- seq data in this study is GEO: GSE191204.

## Author contributions

Conceptualization, M.Y.; Methodology, M.Y. and S.H.; Formal Analysis, M.Y. and S.U.; Investigation, M.Y.; Resource, T.M. and T.O.; Writing – Original Draft, M.Y.; Writing – Review & Editing, S.H., T.I., M.N., and K.Y.; Visualization, M.Y.; Supervision, M.Y., T.I., and K.Y.

## Acknowledgements

We would like to thank M. Yamamoto and M. Muramatsu (Kumamoto University) for insightful discussions, T. Takahashi (Aichi Cancer Center) for useful advices, Daikuzono, Y Ikeda, Y. Fukuchi (Kumamoto University), and S. Fujimura (Liaison Laboratory Research Promotion Center) for technical assistance, and K. Kurokawa (Devers Eye Institute) and S. Watanabe (Kumamoto University) for helpful comments on the manuscript. We would like to thank T. Moroishi (Kumamoto University) for critical reading of this manuscript and providing the HEK293T cells. We also thank S. Okada (Kumamoto University) for providing the *Rag2/Jak3* double-deficient mice. Additionally, we would like to thank T. Takeuchi and H. Nishiyama (Tsukuba Human Tissue Bank Center, University of Tsukuba Hospital) for support in providing the resources. This work was supported by grant-in-aid for Scientific Research (19H03711, 20H03691), by JST SPRING (JPMJSP2127), and by grant from Center for Metabolic Regulation of Health Aging (CMHA).

## Notes

**Declaration of competing interest** The authors declare no competing interests.

### Competing Interest Statement

The authors have declared no competing interest.

### Summary of Updates

Supplemental files updated.

## References

Al-Hajj M, Wicha MS, Benito-Hernandez A, Morrison SJ & Clarke MF (2003) Prospective identification of tumorigenic breast cancer cells. Proc Natl Acad Sci U S A 100: 3983–3988

Awad MM, Liu S, Rybkin II, Arbour KC, Dilly J, Zhu VW, Johnson ML, Heist RS, Patil T, Riely GJ, et al (2021) Acquired Resistance to KRAS G12C Inhibition in Cancer. N Engl J Med 384: 2382–2393

Bailey JM, Alsina J, Rasheed ZA, McAllister FM, Fu YY, Plentz R, Zhang H, Pasricha PJ, Bardeesy N, Matsui W, et al (2014) DCLK1 marks a morphologically distinct subpopulation of cells with stem cell properties in preinvasive pancreatic cancer. Gastroenterology 146: 245–256

Barker N, Ridgway RA, Van Es JH, Van De Wetering M, Begthel H, Van Den Born M, Danenberg E, Clarke AR, Sansom OJ & Clevers H (2009) Crypt stem cells as the cells-of-origin of intestinal cancer. Nature 457: 608–611

Bartucci M, Dattilo R, Moriconi C, Pagliuca A, Mottolese M, Federici G, Di Benedetto A, Todaro M, Stassi G, Sperati F, et al (2015) TAZ is required for metastatic activity and chemoresistance of breast cancer stem cells. Oncogene 34: 681–690

Basu-Roy U, Bayin NS, Rattanakorn K, Han E, Placantonakis DG, Mansukhani A & Basilico C (2015) Sox2 antagonizes the Hippo pathway to maintain stemness in cancer cells. Nat Commun 6: 1–14

Batlle E & Clevers H (2017) Cancer stem cells revisited. Nat Med 23: 1124–1134 doi:10.1038/nm.4409

Bayik D & Lathia JD (2021) Cancer stem cell–immune cell crosstalk in tumour progression. Nat Rev Cancer 21: 526–536 doi:10.1038/s41568-021-00366-w

Bonnet D & Dick JE (1997) Human acute myeloid leukemia is organized as a hierarchy that originates from a primitive hematopoietic cell. Nat Med 3: 730–737

Boumahdi S & de Sauvage FJ (2020) The great escape: tumour cell plasticity in resistance to targeted therapy. Nat Rev Drug Discov 19: 39–56 doi:10.1038/s41573-019-0044-1

Canon J, Rex K, Saiki AY, Mohr C, Cooke K, Bagal D, Gaida K, Holt T, Knutson CG, Koppada N, et al (2019) The clinical KRAS(G12C) inhibitor AMG 510 drives anti-tumour immunity. Nature 575: 217–223

Cao W, Li M, Liu J, Zhang S, Noordam L, Verstegen MMA, Wang L, Ma B, Li S, Wang W, et al (2020) LGR5 marks targetable tumor-initiating cells in mouse liver cancer. Nat Commun 11

Chen YJ, Chen CM, Twu NF, Yen MS, Lai CR, Wu HH, Wang PH & Yuan CC (2009) Overexpression of Aurora B is associated with poor prognosis in epithelial ovarian cancer patients. Virchows Arch 455: 431–440

Choi MY, Widhopf GF, Ghia EM, Kidwell RL, Hasan MK, Yu J, Rassenti LZ, Chen L, Chen Y, Pittman E, et al (2018) Phase I Trial: Cirmtuzumab Inhibits ROR1 Signaling and Stemness Signatures in Patients with Chronic Lymphocytic Leukemia. Cell Stem Cell 22: 951–959.e3

Clarke MF, Dick JE, Dirks PB, Eaves CJ, Jamieson CHM, Jones DL, Visvader J, Weissman IL & Wahl GM (2006) Cancer stem cells - Perspectives on current status and future directions: AACR workshop on cancer stem cells. In Cancer Research pp 9339–9344. American Association for Cancer Research

Corces MR, Trevino AE, Hamilton EG, Greenside PG, Sinnott-Armstrong NA, Vesuna S, Satpathy AT, Rubin AJ, Montine KS, Wu B, et al (2017) An improved ATAC- seq protocol reduces background and enables interrogation of frozen tissues. Nat Methods 14: 959–962

Cordenonsi M, Zanconato F, Azzolin L, Forcato M, Rosato A, Frasson C, Inui M, Montagner M, Parenti AR, Poletti A, et al (2011) The hippo transducer TAZ confers cancer stem cell-related traits on breast cancer cells. Cell 147: 759–772

Cui B, Zhang S, Chen L, Yu J, Widhopf GF, Fecteau JF, Rassenti LZ & Kipps TJ (2013) Targeting ROR1 inhibits epithelial-mesenchymal transition and metastasis. Cancer Res 73: 3649–3660

Daikuzono H, Yamazaki M, Sato Y, Takahashi T & Yamagata K (2021) Development of a DELFIA method to detect oncofetal antigen ROR1-positive exosomes. Biochem Biophys Res Commun 578: 170–176

Dalerba P, Dylla SJ, Park IK, Liu R, Wang X, Cho RW, Hoey T, Gurney A, Huang EH, Simeone DM, et al (2007) Phenotypic characterization of human colorectal cancer stem cells. Proc Natl Acad Sci U S A 104: 10158–10163

Dominguez CX, Müller S, Keerthivasan S, Koeppen H, Hung J, Gierke S, Breart B, Foreman O, Bainbridge TW, Castiglioni A, et al (2020) Single-Cell RNA Sequencing Reveals Stromal Evolution into LRRC15 + Myofibroblasts as a Determinant of Patient Response to Cancer Immunotherapy. Cancer Discov 10: 232–253

Dongre A & Weinberg RA (2019) New insights into the mechanisms of epithelial– mesenchymal transition and implications for cancer. Nat Rev Mol Cell Biol 20: 69– 84

Doroshow DB, Eder JP & LoRusso PM (2017) BET inhibitors: a novel epigenetic approach. Ann Oncol 28: 1776–1787

Ellen TP, Ke Q, Zhang P & Costa M (2008) NDRG1, a growth and cancer related gene: Regulation of gene expression andfunction in normal and disease states. Carcinogenesis 29: 2–8 doi:10.1093/carcin/bgm200

Filippakopoulos P, Qi J, Picaud S, Shen Y, Smith WB, Fedorov O, Morse EM, Keates T, Hickman TT, Felletar I, et al (2010) Selective inhibition of BET bromodomains. Nature 468: 1067–1073

Fox RG, Lytle NK, Jaquish D V., Park FD, Ito T, Bajaj J, Koechlein CS, Zimdahl B, Yano M, Kopp JL, et al (2016) Image-based detection and targeting of therapy resistance in pancreatic adenocarcinoma. Nature 534: 407–411

Frank SB, Schulz V V. & Miranti CK (2017) A streamlined method for the design and cloning of shRNAs into an optimized Dox-inducible lentiviral vector. BMC Biotechnol 17: 1–10

Harvey KF, Zhang X & Thomas DM (2013) The Hippo pathway and human cancer. Nat Rev Cancer 13: 246–257 doi:10.1038/nrc3458

Hayashi H, Higashi T, Yokoyama N, Kaida T, Sakamoto K, Fukushima Y, Ishimoto T, Kuroki H, Nitta H, Hashimoto D, et al (2015) An imbalance in TAZ and YAP expression in hepatocellular carcinoma confers cancer stem cell-like behaviors contributing to disease progression. Cancer Res 75: 4985–4997

Hermann PC, Huber SL, Herrler T, Aicher A, Ellwart JW, Guba M, Bruns CJ & Heeschen C (2007) Distinct Populations of Cancer Stem Cells Determine Tumor Growth and Metastatic Activity in Human Pancreatic Cancer. Cell Stem Cell 1: 313–323

Johnson R & Halder G (2014) The two faces of Hippo: Targeting the Hippo pathway for regenerative medicine and cancer treatment. Nat Rev Drug Discov 13: 63–79 doi:10.1038/nrd4161

Kleeff J, Korc M, Apte M, La Vecchia C, Johnson CD, Biankin A V., Neale RE, Tempero M, Tuveson DA, Hruban RH, et al (2016) Pancreatic cancer. Nat Rev Dis Prim 2: 1–23

Kubota SI, Takahashi K, Nishida J, Tainaka K, Miyazono K & Ueda Correspondence HR (2017) Whole-Body Profiling of Cancer Metastasis with Single-Cell Resolution. Cell Rep 20: 236–250

Langmead B & Salzberg SL (2012) Fast gapped-read alignment with Bowtie 2. Nat Methods 9: 357–359

Lemmon MA & Schlessinger J (2010) Cell signaling by receptor tyrosine kinases. Cell 141: 1117–1134 doi:10.1016/j.cell.2010.06.011

Lens SMA, Voest EE & Medema RH (2010) Shared and separate functions of polo-like kinases and aurora kinases in cancer. Nat Rev Cancer 10: 825–841 doi:10.1038/nrc2964

Li C, Heidt DG, Dalerba P, Burant CF, Zhang L, Adsay V, Wicha M, Clarke MF & Simeone DM (2007) Identification of pancreatic cancer stem cells. Cancer Res 67: 1030–1037

Li C, Wang S, Xing Z, Lin A, Liang K, Song J, Hu Q, Yao J, Chen Z, Park PK, et al (2017) A ROR1-HER3-lncRNA signalling axis modulates the Hippo-YAP pathway to regulate bone metastasis. Nat Cell Biol 19: 106–119

Love MI, Huber W & Anders S (2014) Moderated estimation of fold change and dispersion for RNA-seq data with DESeq2. Genome Biol 15: 1–21

Lytle NK, Ferguson LP, Rajbhandari N, Gilroy K, Fox RG, Deshpande A, Schürch CM, Hamilton M, Robertson N, Lin W, et al (2019) A Multiscale Map of the Stem Cell State in Pancreatic Adenocarcinoma. Cell 177: 572–586.e22

Magee JA, Piskounova E & Morrison SJ (2012) Cancer Stem Cells: Impact, Heterogeneity, and Uncertainty. Cancer Cell 21: 283–296

La Manno G, Soldatov R, Zeisel A, Braun E, Hochgerner H, Petukhov V, Lidschreiber K, Kastriti ME, Lönnerberg P, Furlan A, et al (2018) RNA velocity of single cells. Nature 560: 494–498

Mao Z, Xiao H, Shen P, Yang Y, Xue J, Yang Y, Shang Y, Zhang L, Li X, Zhang Y, et al (2022) KRAS(G12D) can be targeted by potent inhibitors via formation of salt bridge. Cell Discov 8: 1–14

Marusyk A, Janiszewska M & Polyak K (2020) Intratumor Heterogeneity: The Rosetta Stone of Therapy Resistance. Cancer Cell 37: 471–484 doi:10.1016/j.ccell.2020.03.007

Mizrahi JD, Surana R, Valle JW & Shroff RT (2020) Pancreatic cancer. Lancet 395: 2008–2020

O’Brien CA, Pollett A, Gallinger S & Dick JE (2007) A human colon cancer cell capable of initiating tumour growth in immunodeficient mice. Nature 445: 106– 110

Okada S, Ono A, Hattori S, Kariya R, Iwanaga S, Taura M, Harada H & Suzu S (2011) Comparative study of human hematopoietic cell engraftment into Balb/c and C57BL/6 strain of rag-2/Jak3 double-deficient mice. J Biomed Biotechnol 2011

Oki S, Ohta T, Shioi G, Hatanaka H, Ogasawara O, Okuda Y, Kawaji H, Nakaki R, Sese J & Meno C (2018) Ch IP -Atlas: a data-mining suite powered by full integration of public Ch IP -seq data. EMBO Rep 19: e46255

Osawa H, Takahashi H, Nishimura J, Ohta K, Haraguchi N, Hata T, Yamamoto H, Mizushima T, Takemasa I, Doki Y, et al (2016) Full-length LGR5-positive cells have chemoresistant characteristics in colorectal cancer. Br J Cancer 114: 1251– 1260

Pastushenko I, Brisebarre A, Sifrim A, Fioramonti M, Revenco T, Boumahdi S, Van Keymeulen A, Brown D, Moers V, Lemaire S, et al (2018) Identification of the tumour transition states occurring during EMT. Nature 556: 463–468

Pastushenko I, Mauri F, Song Y, de Cock F, Meeusen B, Swedlund B, Impens F, Van Haver D, Opitz M, Thery M, et al (2021) Fat1 deletion promotes hybrid EMT state, tumour stemness and metastasis. Nature 589: 448–455

Patro R, Duggal G, Love MI, Irizarry RA & Kingsford C (2017) Salmon provides fast and bias-aware quantification of transcript expression. Nat Methods 14: 417–419

Puram S V., Tirosh I, Parikh AS, Patel AP, Yizhak K, Gillespie S, Rodman C, Luo CL, Mroz EA, Emerick KS, et al (2017) Single-Cell Transcriptomic Analysis of Primary and Metastatic Tumor Ecosystems in Head and Neck Cancer. Cell 171: 1611–1624.e24

Ramírez F, Ryan DP, Grüning B, Bhardwaj V, Kilpert F, Richter AS, Heyne S, Dündar F & Manke T (2016) deepTools2: a next generation web server for deep- sequencing data analysis. Nucleic Acids Res 44: W160–W165

Raphael BJ, Hruban RH, Aguirre AJ, Moffitt RA, Yeh JJ, Stewart C, Robertson AG, Cherniack AD, Gupta M, Getz G, et al (2017) Integrated Genomic Characterization of Pancreatic Ductal Adenocarcinoma. Cancer Cell 32: 185–203.e13

Reid MD, Basturk O, Thirabanjasak D, Hruban RH, Klimstra DS, Bagci P, Altinel D & Adsay V (2011) Tumor-infiltrating neutrophils in pancreatic neoplasia. Mod Pathol 24: 1612–1619

Ricci-Vitiani L, Lombardi DG, Pilozzi E, Biffoni M, Todaro M, Peschle C & De Maria R (2007) Identification and expansion of human colon-cancer-initiating cells. Nature 445: 111–115

Seino T, Kawasaki S, Shimokawa M, Tamagawa H, Toshimitsu K, Fujii M, Ohta Y, Matano M, Nanki K, Kawasaki K, et al (2018) Human Pancreatic Tumor Organoids Reveal Loss of Stem Cell Niche Factor Dependence during Disease Progression. Cell Stem Cell 22: 454–467.e6

Shackleton M, Quintana E, Fearon ER & Morrison SJ (2009) Heterogeneity in Cancer: Cancer Stem Cells versus Clonal Evolution. Cell 138: 822–829

Shibue T & Weinberg RA (2017) EMT, CSCs, and drug resistance: The mechanistic link and clinical implications. Nat Rev Clin Oncol 14: 611–629 doi:10.1038/nrclinonc.2017.44

Shimokawa M, Ohta Y, Nishikori S, Matano M, Takano A, Fujii M, Date S, Sugimoto S, Kanai T & Sato T (2017) Visualization and targeting of LGR5 + human colon cancer stem cells. Nature 545: 187–192

Shimomura O, Oda T, Tateno H, Ozawa Y, Kimura S, Sakashita S, Noguchi M, Hirabayashi J, Asashima M & Ohkohchi N (2018) A Novel Therapeutic Strategy for Pancreatic Cancer: Targeting Cell Surface Glycan Using rBC2LC-N Lectin– Drug Conjugate (LDC). Mol Cancer Ther 17: 183–195

Singh SK, Hawkins C, Clarke ID, Squire JA, Bayani J, Hide T, Henkelman RM, Cusimano MD, Dirks PB, Terasaki M, et al (2003) Identification of a cancer stem cell in human brain tumors. Cancer Res 63: 5821–8

Skene PJ & Henikoff S (2017) An efficient targeted nuclease strategy for high-resolution mapping of DNA binding sites. Elife 6

Song S, Ajani JA, Honjo S, Maru DM, Chen Q, Scott AW, Heallen TR, Xiao L, Hofstetter WL, Weston B, et al (2014) Hippo coactivator YAP1 upregulates SOX9 and endows esophageal Cancer cells with stem-like properties. Cancer Res 74: 4170–4182

De Sousa E Melo F, Kurtova A V., Harnoss JM, Kljavin N, Hoeck JD, Hung J, Anderson JE, Storm EE, Modrusan Z, Koeppen H, et al (2017) A distinct role for Lgr5 + stem cells in primary and metastatic colon cancer. Nature 543: 676–680

Srivastava S, Furlan SN, Jaeger-Ruckstuhl CA, Sarvothama M, Berger C, Smythe KS, Garrison SM, Specht JM, Lee SM, Amezquita RA, et al (2021) Immunogenic Chemotherapy Enhances Recruitment of CAR-T Cells to Lung Tumors and Improves Antitumor Efficacy when Combined with Checkpoint Blockade. Cancer Cell 39: 193–208.e10

Srivastava S, Salter AI, Liggitt D, Yechan-Gunja S, Sarvothama M, Cooper K, Smythe KS, Dudakov JA, Pierce RH, Rader C, et al (2019) Logic-Gated ROR1 Chimeric Antigen Receptor Expression Rescues T Cell-Mediated Toxicity to Normal Tissues and Enables Selective Tumor Targeting. Cancer Cell 35: 489–503.e8

Stuart T, Butler A, Hoffman P, Hafemeister C, Papalexi E, Mauck WM, Hao Y, Stoeckius M, Smibert P & Satija R (2019) Comprehensive Integration of Single- Cell Data. Cell 177: 1888–1902.e21

Subramanian A, Tamayo P, Mootha VK, Mukherjee S, Ebert BL, Gillette MA, Paulovich A, Pomeroy SL, Golub TR, Lander ES, et al (2005) Gene set enrichment analysis: A knowledge-based approach for interpreting genome-wide expression profiles. Proc Natl Acad Sci U S A 102: 15545–15550

Vischioni B, Oudejans JJ, Vos W, Rodriguez JA & Giaccone G (2006) Frequent overexpression of aurora B kinase, a novel drug target, in non-small cell lung carcinoma patients. Mol Cancer Ther 5: 2905–2918

Wang X, Allen S, Blake JF, Bowcut V, Briere DM, Calinisan A, Dahlke JR, Fell JB, Fischer JP, Gunn RJ, et al (2022) Identification of MRTX1133, a Noncovalent, Potent, and Selective KRASG12DInhibitor. J Med Chem 65: 3123–3133

Xue JY, Zhao Y, Aronowitz J, Mai TT, Vides A, Qeriqi B, Kim D, Li C, de Stanchina E, Mazutis L, et al (2020) Rapid non-uniform adaptation to conformation-specific KRAS(G12C) inhibition. Nature 577: 421–425

Yamaguchi T, Yanagisawa K, Sugiyama R, Hosono Y, Shimada Y, Arima C, Kato S, Tomida S, Suzuki M, Osada H, et al (2012) NKX2-1/TITF1/TTF-1-Induced ROR1 Is Required to Sustain EGFR Survival Signaling in Lung Adenocarcinoma. Cancer Cell 21: 348–361

Zeng L & Zhou M-M (2002) Bromodomain: an acetyl-lysine binding domain. FEBS Lett 513: 124–128

Zeng WF, Navaratne K, Prayson RA & Weil RJ (2007) Aurora B expression correlates with aggressive behaviour in glioblastoma multiforme. J Clin Pathol 60: 218

Zhang S, Chen L, Cui B, Chuang HY, Yu J, Wang-Rodriguez J, Tang L, Chen G, Basak GW & Kipps TJ (2012a) ROR1 is expressed in human breast cancer and associated with enhanced tumor-cell growth. PLoS One 7: 31127

Zhang S, Chen L, Wang-Rodriguez J, Zhang L, Cui B, Frankel W, Wu R & Kipps TJ (2012b) The onco-embryonic antigen ROR1 is expressed by a variety of human cancers. Am J Pathol 181: 1903–1910

Zhang S, Cui B, Lai H, Liu G, Ghia EM, Widhopf GF, Zhang Z, Wu CCN, Chen L, Wu R, et al (2014) Ovarian cancer stem cells express ROR1, which can be targeted for anti-cancer-stem-cell therapy. Proc Natl Acad Sci U S A 111: 17266–17271

Zhang S, Zhang H, Ghia EM, Huang J, Wu L, Zhang J, Lam S, Lei Y, He J, Cui B, et al (2019) Inhibition of chemotherapy resistant breast cancer stem cells by a ROR1 specific antibody. Proc Natl Acad Sci U S A 116: 1370–1377

Zhang Y, Liu T, Meyer CA, Eeckhoute J, Johnson DS, Bernstein BE, Nussbaum C, Myers RM, Brown M, Li W, et al (2008) Model-based analysis of ChIP-Seq (MACS). Genome Biol 9: 1–9

Zhu Q, Liu N, Orkin SH & Yuan GC (2019) CUT and RUNTools: A flexible pipeline for CUT and RUN processing and footprint analysis. Genome Biol 20: 1–12

