## Supplemental Figure for "ROR1 plays a critical role in pancreatic tumor-initiating cells with a partial EMT signature"

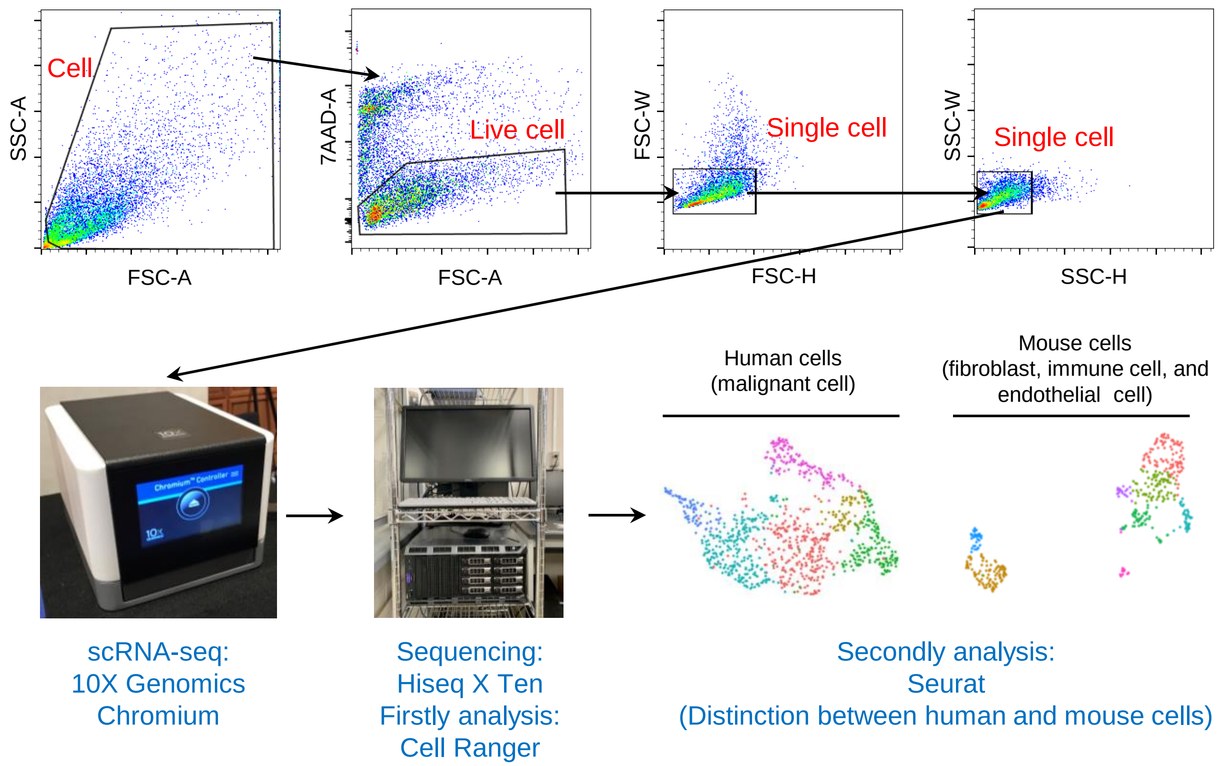


**Supplemental Figure 1. Related to Figure 1. Schematic diagram for the generation of single-cell RNA sequencing data.**

Representative FACS plots of the S2-VP10 xenograft are shown. The sorted population did not include dead and duplicate cells. Human and mouse cells were distinguished by data analysis using Cell Ranger and Seurat.

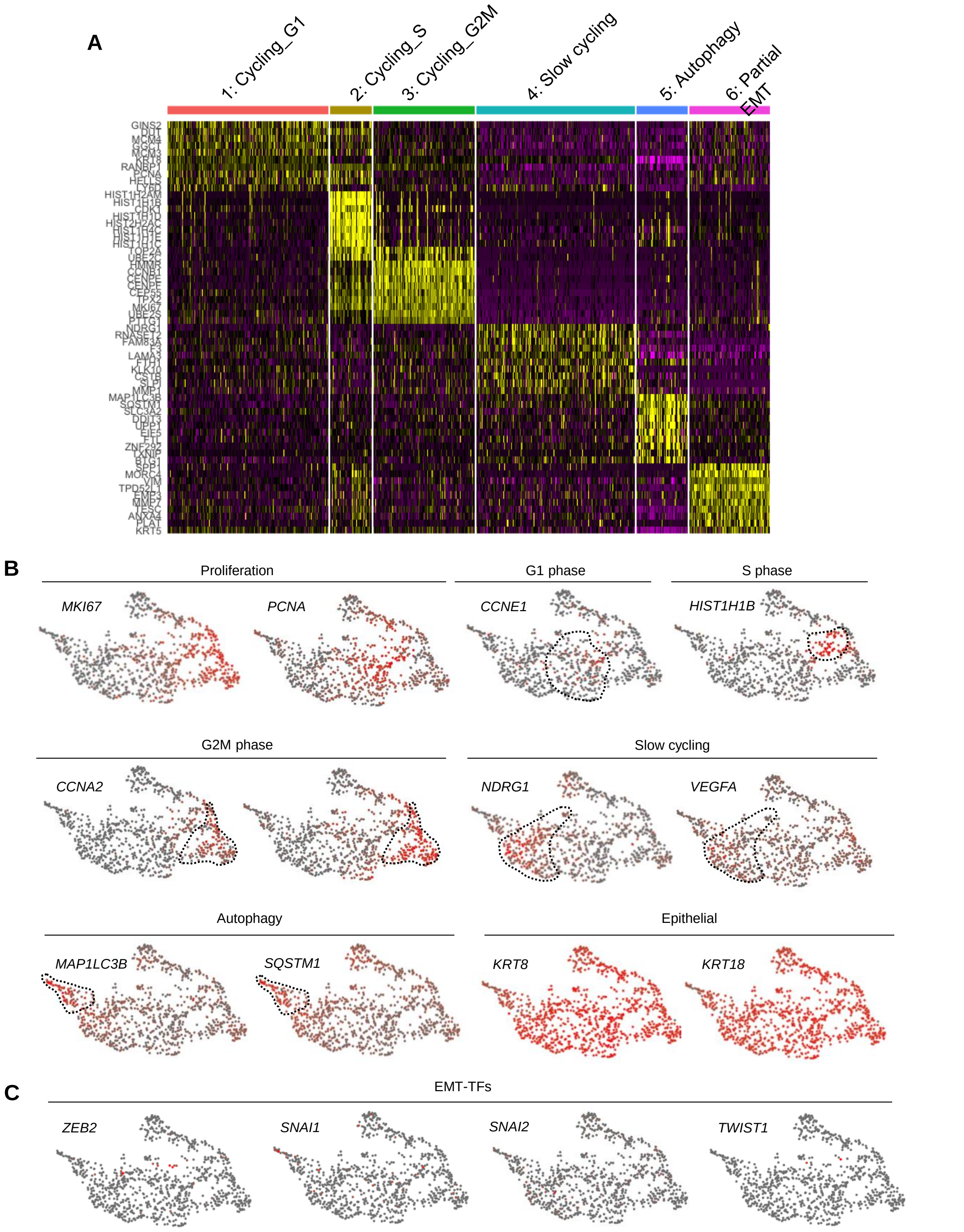


**Supplemental Figure 2. Related to Figure 1. scRNA-seq data in S2-VP10 xenografts.**
**(A)** Heatmap showing the expression of cluster-specific genes.
**(B)–(D)** UMAP plots displaying selected marker gene expression.

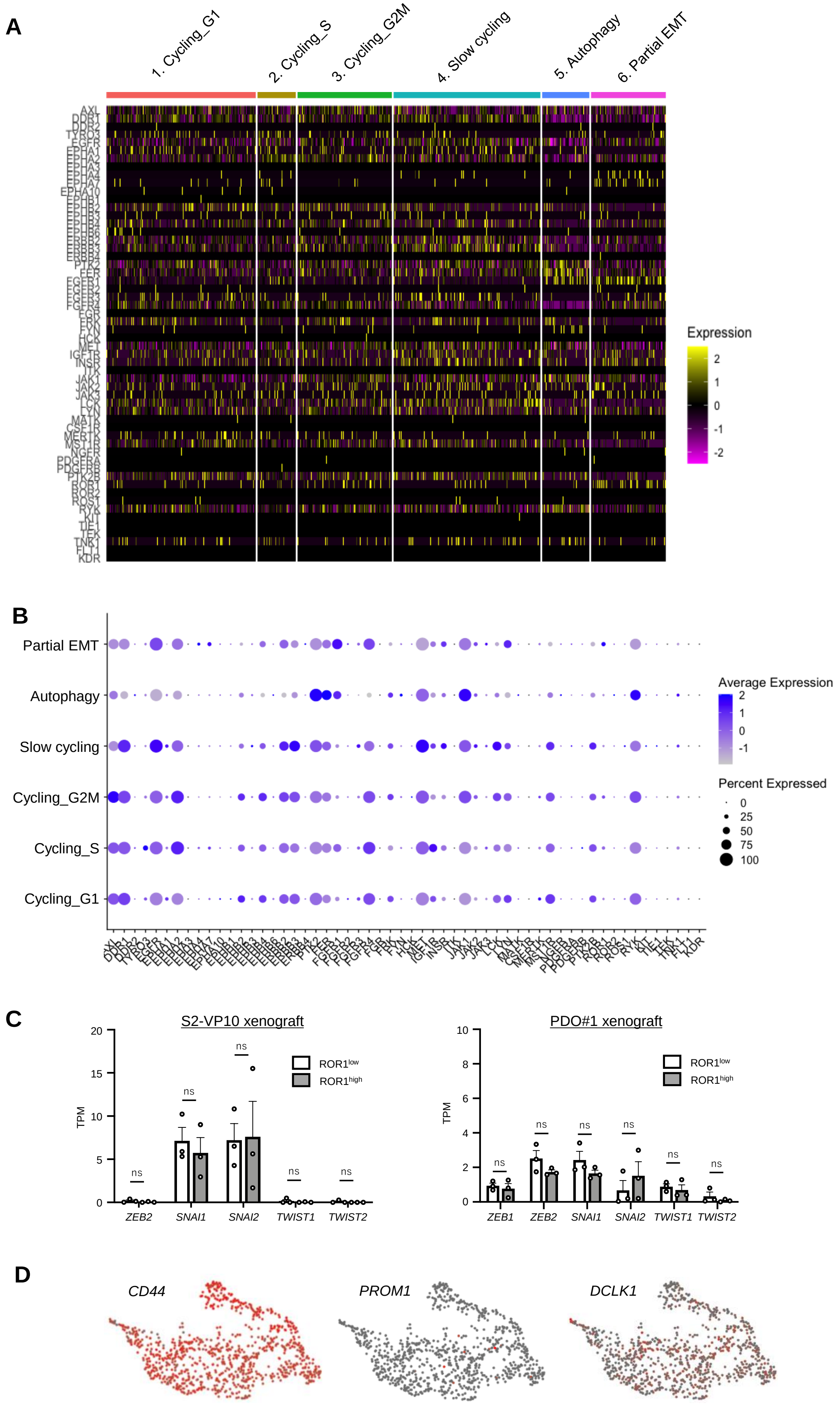


Supplemental Figure 3

**Supplemental Figure 3. Related to Figure 2. Identification of the markers for partial EMT populations in PDAC.**

**(A)** Heatmap showing the expression of RTK genes within individual cells.

**(B)** Dot plot displaying the expression of RTK genes in subpopulations.

**(C)** Expression levels of classical EMT TF genes in $ROR1^{high}$ cells and $ROR1^{low}$ cells from S2-VP10 and PDO#1 xenografts (n = 3).

**(D)** UMAP plots displaying known CSC marker gene expression.

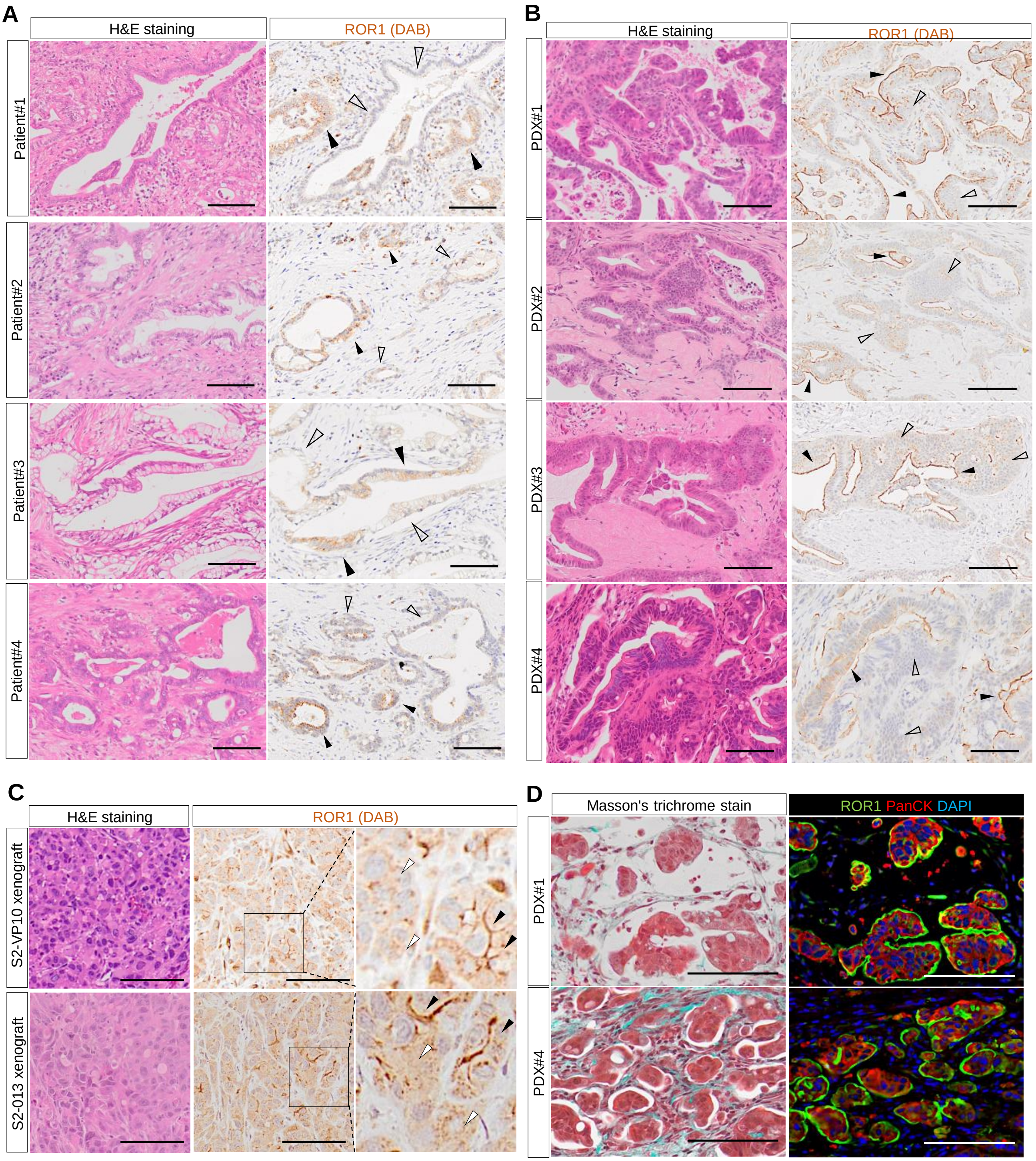


**Supplemental Figure 4. Related to Figure 3. ROR1 immunostaining in patient tumors and xenografts.** **(A)–(C)** Representative images of patient PDAC (A), parental PDXs (B), and cell-derived xenografts (from S2-VP10 and S2-013) (C) stained with H&E and for ROR1. Black arrow, positive areas; white arrow, negative areas of ROR1. **(D)** Representative images of Masson's trichrome and immunohistochemical staining for ROR1 and PanCK in patient-derived xenografts (PDX).

Scale bars, 100 μm (A–D).

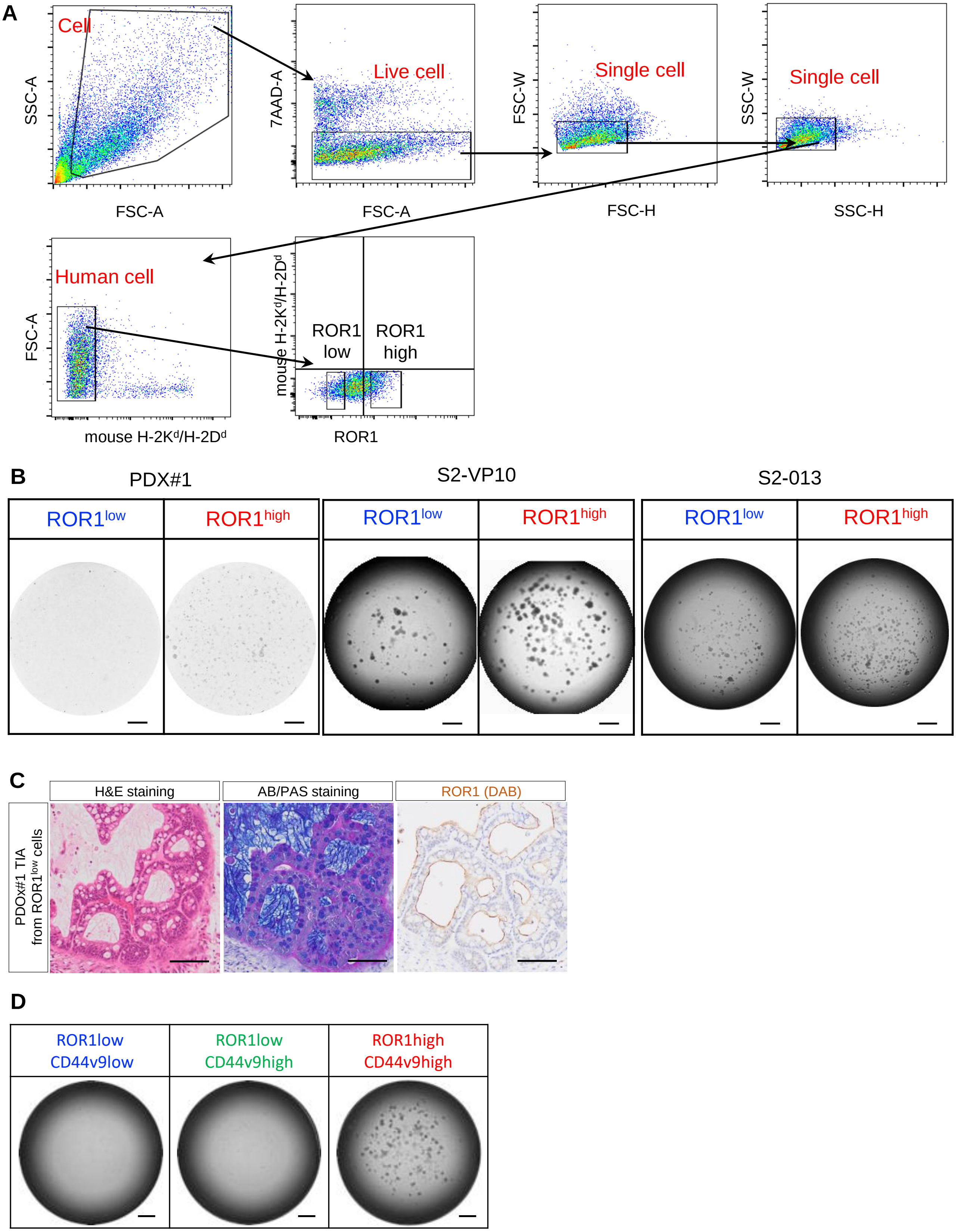




Supplemental Figure 5

**Supplemental Figure 5. Related to Figure 3. Tumorigenicity of isolated intratumor ROR1$^{high}$ cells.**
**(A)** Gating scheme for the isolation of ROR1$^{high}$ and ROR1$^{low}$ cells from xenografts. The sorted population contained only single, viable human cancer cells.
**(B)** Representative images of organoid or colony formation assays. The ROR1$^{high}$ cells from PDX#1 and xenografts (S2-VP10, S2-013) efficiently formed organoids or colonies compared with ROR1$^{low}$ cells.
**(C)** Representative images stained with H&E and AB/PAS, and for ROR1 in tumor derived from ROR1$^{low}$ cells of PDO#1 xenograft. ROR1 was again expressed heterogeneously.
**(D)** Representative images of colony formation assays. The ROR1$^{high}$/CD44v9$^{high}$ cells from S2-VP10 xenografts efficiently formed colonies.

Scale bars, 1 mm (b and d) and 100 µm (c).

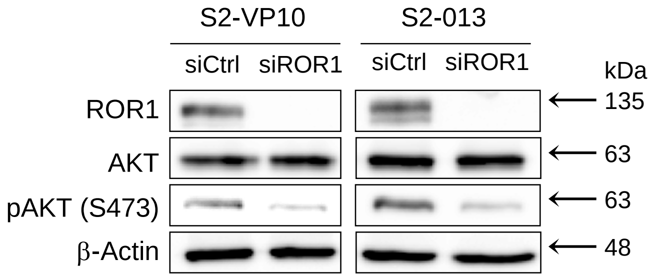


**Supplemental Figure 6. Related to Figure 4. Western blot analysis of ROR1, AKT, phospho-AKT (Ser473), and β-actin in S2-VP10 and S2-013 cells transfected with control or ROR1 siRNA.**

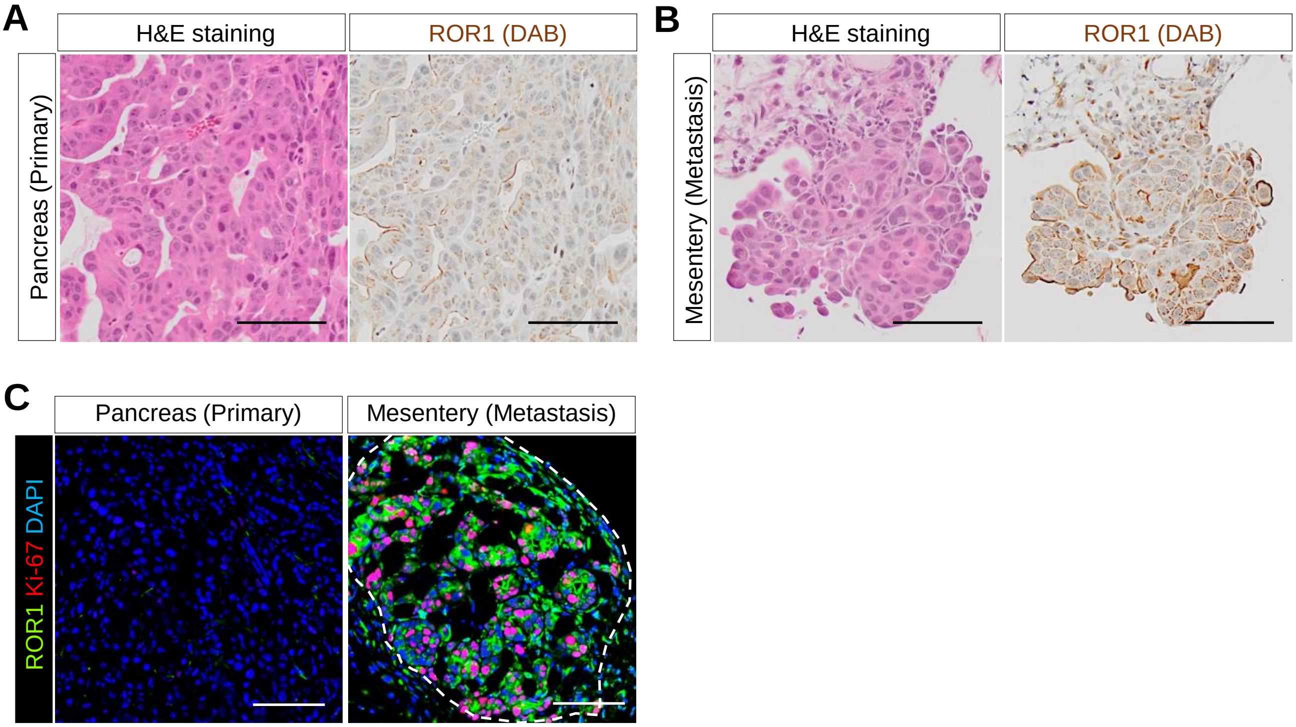


**Supplemental Figure 7. Related to Figure 6. Histological analysis in primary and metastasis sites of S2-013 xenografts.**
**(A), (B)** Representative images of S2-013 xenografts stained with H&E (A) and for ROR1 (B).
**(C)** Representative images of S2-013 xenografts co-stained for ROR1 and Ki-67. The dotted line indicates the metastatic lesions.

Scale bars, 100 µm (A–C)

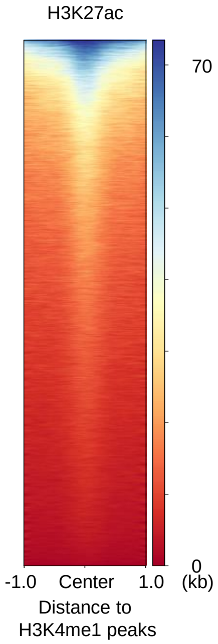


**Supplemental Figure 8. Related to Figure 8. Heatmap of H3K27ac CUT&RUN data on the genomic locations of the H3K4me1 peaks in S2-VP10 cells.**

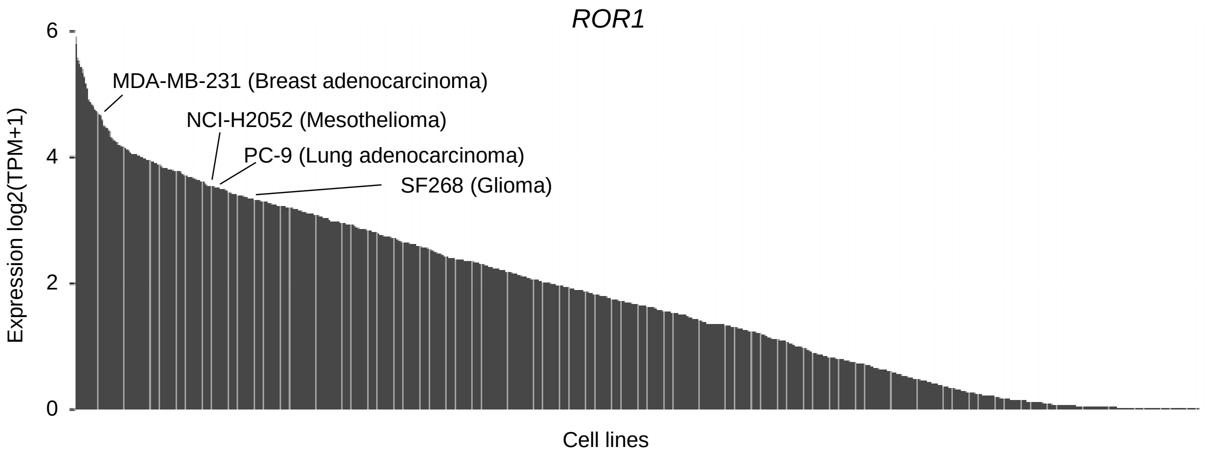


**Supplemental Figure 9. Related to Figure 8. Bar plot of ROR1 expression in cell lines from the CCLE database.** Bar plot showing ROR1 expression of 1389 cell lines.
